# Evolution of parent-of-origin effects on placental gene expression in house mice

**DOI:** 10.1101/2023.08.24.554674

**Authors:** Fernando Rodriguez-Caro, Emily C. Moore, Jeffrey M. Good

## Abstract

The mammalian placenta is a hotspot for the evolution of genomic imprinting, a form of gene regulation that involves the parent-specific epigenetic silencing of one allele. Imprinted genes are central to placental development and are thought to contribute to the evolution of reproductive barriers between species. However, it is unclear how rapidly imprinting evolves or how functional specialization among placental tissues influences the evolution of imprinted expression. We compared parent-of-origin expression bias across functionally distinct placental layers sampled from reciprocal crosses within three closely related lineages of mice (*Mus*). Using genome-wide gene expression and DNA methylation data from fetal and maternal tissues, we developed an analytical strategy to minimize pervasive bias introduced by maternal contamination of placenta samples. We corroborated imprinted expression at 42 known imprinted genes and identified five candidate imprinted genes showing parent-of-origin specific expression and DNA methylation. Paternally-biased expression was enriched in the labyrinth zone, a layer specialized in nutrient transfer, and maternally-biased genes were enriched in the junctional zone, which specializes in modulation of maternal physiology. Differentially methylated regions were predominantly determined through epigenetic modification of the maternal genome and were associated with both maternally- and paternally-biased gene expression. Lastly, comparisons between lineages revealed a small set of co-regulated genes showing rapid divergence in expression levels and imprinted status in the *M. m. domesticus* lineage. Together, our results reveal important links between core functional elements of placental biology and the evolution of imprinted gene expression among closely related rodent species.

## Introduction

Embryonic development in eutherian mammals relies on the placenta, a transitory organ that mediates physiological exchange and modulates maternal metabolism during pregnancy. Although placentation has evolved in several vertebrate lineages (Stewart and Thompson 2000; Wildman *et al*. 2006; Furness *et al*. 2021), the mammalian placenta exhibits a remarkable level of specialization reflected in a diverse array of specialized cells and a high degree of maternal-fetal tissue integration (Blackburn 2015; Roberts *et al*. 2016). This specialization was achieved through the emergence and diversification of the trophoblast cell lineage (Selwood and Johnson 2006) and accompanied by the evolution of a unique form of gene regulation known as genomic imprinting, an epigenetic phenomena whereby certain genes show parent-of-origin dependent allelic transcription bias. Imprinted expression is a critical element of placental development and is thought to evolve under strong selective pressures (Moore and Haig 1991). However, despite its importance for mammalian development and evolution, our understanding of natural variation of imprinted expression across placental tissues and between species remains limited.

Despite considerable morphological diversity across mammalian taxa (Roberts *et al*. 2016), the functional architecture of the placenta is broadly conserved across species (Cross *et al*. 2003). In most mammals, the placenta is compartmentalized into two distinct functional layers, known as the labyrinth and junctional zones in rodents (Figure 1A). The labyrinth zone contains fetal and maternal capillaries that irrigate the syncytiotrophoblast, a tissue composed of multinucleated cells that is in direct contact with maternal blood and mediates nutrient exchange (Simmons *et al*. 2008). The junctional zone serves endocrine functions and is composed of invasive trophoblast cells largely involved in the production of signaling molecules that mediate maternal-fetal crosstalk during pregnancy (Simmons *et al*. 2008; Hu and Cross 2010). In lineages with invasive placentation (*e*.*g*., rodents and primates), cells derived from the internal lining of the uterine wall form a third placental layer known as the maternal decidua (Abrahamsohn and Zorn 1993). The decidua plays a critical role in the modulation of the maternal immune response, creating a local uterine environment that prevents rejection of embryonic cells during pregnancy

**Figure 1.**
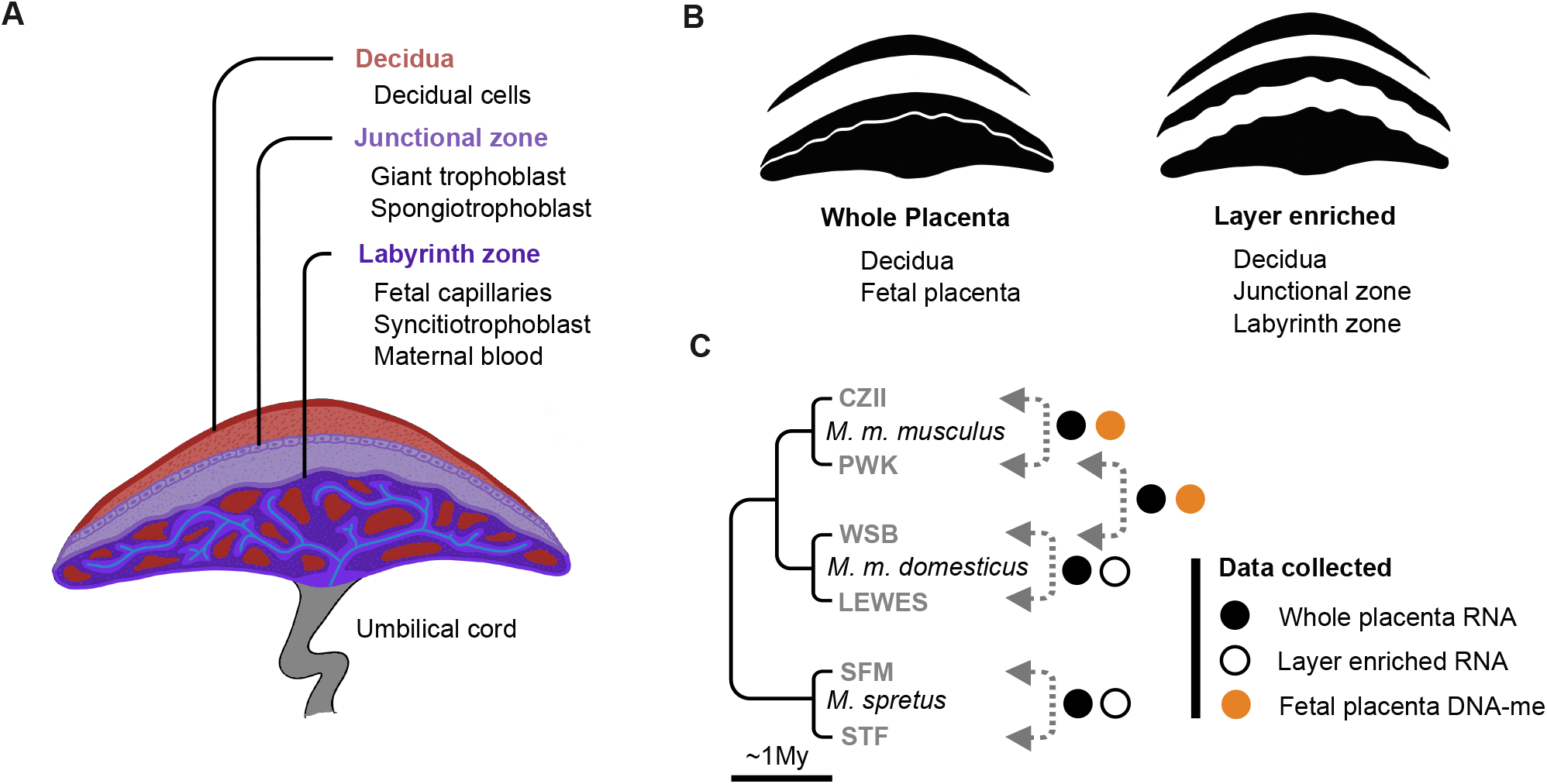
Experimental design. **A)** Schematic representation of placental anatomy and maternal-fetal tissue integration. The three major layers in the mouse placenta include the decidua, of maternal origin, and the junctional and labyrinth zones, of embryonic origin. **B)** Types of placenta dissections performed for this experiment. Whole placenta dissections include the separation of maternal and fetal tissues, while layer enriched dissections include further separation of the fetal placenta into its two major compartments. **C)** Phylogenetic relations among the three mouse lineages included in this study and data collected from each type of reciprocal cross. See Figure S1 for further sampling details.

Successful development and function of the placenta depend on proper regulation of several genes showing parent-of-origin dependent expression often associated with epigenetic marks established during gametogenesis (Kelsey and Feil 2013; Thamban *et al*. 2020). These gametic imprints persist after fertilization at imprinting control regions (ICRs) which influence allele specific silencing at neighboring loci through diverse epigenetic mechanisms (*e*.*g*., somatic DNA-methylation, chromatin modifications, expression of regulatory RNAs, etc.) (Andergassen *et al*. 2021), often forming clusters of imprinted expression in the genome. Most known ICRs originate from oocyte- or sperm-specific DNA methylation marks, known as canonical imprints, but a small set of ICRs are determined by oocyte-specific histone modifications (H3K27me) and these are considered non-canonical imprints (Hanna and Kelsey 2021; Inoue *et al*. 2017b). Canonical imprints are widespread in eutherian mammals and a subset of canonically imprinted genes show conserved imprinted status in all therian species studied (Wang *et al*. 2013; Babak *et al*. 2015; Brekke *et al*. 2016; Richard Albert *et al*. 2023; Cao *et al*. 2023). In contrast, non-canonical imprints have been only documented in rodents and linked to the regulation of ∼16 placenta-specific imprinted genes (Richard Albert *et al*. 2023; Inoue *et al*. 2017b,a). Collectively, these findings suggest considerable variation in the mechanistic underpinnings of imprinting in mammals. However, the evolution of of imprinted expression across functional elements of the placenta and among closely-related species remains poorly understood.

Detailed characterization of placental imprinted expression has been limited to humans and a small set of other distantly related species of agricultural (*e*.*g*., cattle, horses) or biomedical (*e*.*g*., mice, rats) importance (Morison and Reeve 1998; Wei *et al*. 2014). Most of the available data derive from genome-wide allele-specific RNA-seq experiments in known pedigrees or reciprocal crosses between breeds or strains across a small set of distantly related species (Wang *et al*. 2011, 2013; Brekke *et al*. 2016; Pilvar *et al*. 2019; Richard Albert *et al*. 2023). However, detection of imprinted genes based on allelic imbalance in placental tissues has been plagued by maternal tissue contamination in species with invasive placentation (Wang *et al*. 2011; Okae *et al*. 2012; Finn *et al*. 2014), resulting in high false positives rates and low reproducibility across studies (Edwards *et al*. 2023). These limitations have obscured comparisons of the placental imprinted landscape between species (Babak *et al*. 2015). Furthermore, despite the diversity of placental functions and cell types (Pavličev *et al*. 2017), it remains unclear if and how tissue specialization in the mammalian placenta influences the evolution of genomic imprinting. Specialized placental layers evolve under distinct evolutionary pressures (*e*.*g*., (Sandovici *et al*. 2022; Suhail *et al*. 2022; Wilsterman *et al*. 2023) and show differences in the extent of inter-specific epigenetic divergence (Decato *et al*. 2017). Thus, specialization of placental tissues may influence regulatory divergence and select for different patterns of genomic imprinting across functional layers.

Mice (genus *Mus*) provide a powerful system for the study of placental biology and imprinted expression evolution, with extensive intra- and inter-specific variation (Corbet and Hill 1994) and extensive genomic resources, including a high quality reference genome of *Mus musculus* (Mouse Genome Sequencing Consortium *et al*. 2002; Bayona-Bafaluy *et al*. 2003; Church *et al*. 2009). Additionally, several wild derived inbred strains have been established and extensively genotyped, providing a unique opportunity to leverage the genetic diversity of this genus (Keane *et al*. 2011; Nikolskiy *et al*. 2015; Srivastava *et al*. 2017; Chang *et al*. 2017). *Mus musculus* has three subspecies - *M. m. musculus, M. m. domesticus*, and *M. m. castaneus* – which arose within 350-500K years (Salcedo *et al*. 2007; Agwamba and Nachman 2023; Phifer-Rixey *et al*. 2020). Previous studies have used crosses restricted to these subspecies to detect imprinted expression in the whole fetal placenta Wang *et al*. (2011); Okae *et al*. (2012); Finn *et al*. (2014); Andergassen *et al*. (2021), providing an important reference of the placental imprintome, but limiting our understanding of how imprinting evolves between species and across tissue types. Here we build on this foundation by collecting genome-wide gene expression and DNA methylation data from functionally distinct fetal and maternal placental tissues sampled from two of these subspecies (*M. m. musculus* and *M. m. domesticus*) and the closely related species *M. spretus* (Figures 1B,C). Analyzing these data with a novel strategy that minimizes the effect of maternal contamination, we reveal important links between core functional elements of placental biology and the evolution of imprinted gene expression.

## Methods

### Sampling and experimental design

We used six wild-derived inbred strains sampled from two species of mice, *Mus musculus* and *Mus spretus*, spanning ∼1 million years to a most recent common ancestor (Suzuki *et al*. 2004). *Mus musculus* was represented by two wild-derived inbred strains of *M. m. musculus* (PWK/Phj and CZECHII/Eij, hereafter: mus^PWK^ and mus^CZII^) and two wild-derived inbred strains of *M. m. domesticus* (WSB/EiJ and LEWES/EiJ, hereafter: dom^WSB^ and dom^LEW^). *Mus spretus* was represented by two wild-derived strains (STF/Pas and SFM/M, hereafter: spret^STF^ and spret^SFM^) that have been maintained in a closed colony since establishment in 1982. All animals used in this study were bred at the University of Montana using mice purchased from the Jackson Laboratory in 2010 (*M. musculus*) and from the Montpellier Wild Mice Genetic Repository (*M. spretus*). All experimental procedures were performed in accordance with University of Montana Institutional Animal Care and Use Committee regulations (IACUC protocol #068-21).

We performed reciprocal crosses between strains within each of the three lineages (mus^PWK^ x mus^CZII^, dom^WSB^ x dom^LEW^, spret^STF^ x spret^SFM^) and one additional subspecific cross between *M. m. musculus* and *M. m. domesticus* (mus^PWK^ x dom^WSB^) to increase our power to quantify allele-specific expression. Since these wild-derived strains rarely have visible copulatory plugs, female mice were left with males until pregnancy was confirmed, and weighed at mating, at 14 days post mating, and every two days following. Weight gain was used in conjunction with visible cues such as nipple prominence and relative girth to determine onset and stage of pregnancy, determined for each strain through similar measurements of breeding mice taken during the course of colony maintenance. Females were sacrificed in late-stage gestation and embryonic development was determined via Theiler stage (TS 24-26; Theiler 1989).

We collected two types of placental samples from these crosses. For fetal placenta dissections, maternal decidua was separated from fetal placenta (1B, left) and each embryo and fetal placenta were weighed. Fetal placental samples were immediately sectioned into four even quadrants along the sagittal and transverse axes (*i*.*e*., preserving the layer architecture within each section). For layer-enriched dissections (1B, right), maternal decidua tissue was removed and the junctional zone was carefully separated from the labyrinth zone using fine dissection tools (Qu *et al*. 2014). All tissues dissected were snap-frozen in liquid nitrogen. Each pup and its correspondent placenta were sexed through embryo genotyping (Tunster 2017). We selected six samples from layer enriched dissections of two cross types (dom^WSB^ x dom^LEW^, spret^SFM^ x spret^STF^) to quantify layer-specific expression from the two focal species. For each of the four reciprocal crosses, we selected 20 fetal placenta samples for RNA sequencing (five per sex per cross direction). From this larger RNA experiment, we chose four samples each from reciprocal *Mus* (mus^PWK^ x mus^CZII^) and hybrid (mus^PWK^ x dom^WSB^) crosses for whole genome bisulfite sequencing (WGBS). Additionally, we kept the maternal decidua from eight fetal placenta samples per cross type (two per sex per cross direction) for RNA sequencing to generate genotype-specific estimates of maternal to fetal expression ratios.

### Library Preparation

Placental tissue was homogenized in TriReagent RT (Molecular Research Center, cat. no. RT111) using a Qiagen TissueLyser (cat. no. 85300), and RNA was extracted using a hybrid TriReagent – RNeasy spin column method. Following TriReagent phase separation, the aqueous phase was used as input to an RNeasy Mini column (Qiagen, cat. no. 74106), after which the manufacturer’s protocol was followed. Once all tissue collections were complete, we prepared RNA-seq sequencing libraries in blocks containing an equal representation of all cross-types to minimize batch effects. Libraries were generated using the Kapa mRNA HyperPrep kit (Roche, cat. no. KK8581), barcoded with unique dual index adaptors (Roche, cat. no. KK8727), and pooled across two lanes of 150 bp PE Illumina NovaSeqS4 (Novogene, Sacramento, CA). For samples also prepared for WGBS, high quality DNA was extracted from flash frozen placental tissue using a DNeasy Blood and Tissue Kit (Qiagen, cat. no. 69506), and bisulfite converted with the Zymo EZ DNA Methylation Gold spin column kit (cat. no. D5005). Libraries were prepared with the transposase-based Accel-NGS Methyl-Seq DNA Library kit (Swift Biosciences, cat. no. 30096) and barcoded with the Methyl-seq set A indexing kit (Swift Biosciences, cat. no. 36024), following manufacturer’s instructions. WGBS sequencing was performed on a single lane of 150 bp PE Illumina NovaSeqS4 and sequenced by Novogene, (Sacramento, CA).

### Transcriptome data processing

RNA-seq reads were filtered and trimmed using *Trimmomatic* (Bolger *et al*. 2014), with the following settings [ILLUMINACLIP:TruSeq3-PE.fa:2:30:10 LEADING:3 TRAILING:3 SLIDINGWINDOW:5:20 MINLEN = 36]. To reduce reference bias, strain-specific pseudo-genomes were downloaded from the Mouse Collaborative Cross database (WSB, LEWES) or generated with *Modtools* (Huang *et al*. 2014) using resequenced data mapped to the *M. musculus* (PWK, CZII) or *M. spretus* (STF, SFM) reference genome assemblies (GRCm38 and SPRET_EiJ_v1 respectively). Trimmed RNA-seq reads from each sample were mapped to parental pseudogenomes using *Hisat2* (Kim *et al*. 2019). A custom python script was used to reformat *Hisat2* alignment files to allow compatibility with the *ModTools* pipeline (https://github.com/LF-Rodriguez/Mouse_imprinting_evolution). Genome mapping coordinates were then converted to a common reference coordinate system and merged into a single alignment file using *Lapels* (Huang *et al*. 2014). All reads were assigned maternal, paternal, or undefined origin based on differences in mapping quality to each parental pseudogenome using *Suspenders* (Huang *et al*. 2014). Two types of expression matrices were generated. First, 4 Evolution of parent-of-origin effects on placental gene expression all alignments from each sample (regardless of their parent of origin) were used as input into *featureCounts* (Liao *et al*. 2014) to generate gene expression matrices. Then, reads unambiguously assigned to each parent were grouped into separate alignment files and used as input to *featureCounts* to generate allelic expression matrices.

### Layer-induced expression analysis

To detect layer-induced expression, we first categorized genes as either expressed or not expressed in each layer and then identified genes with expression enriched for a particular layer (*i*.*e*., induced expression) following Kopania *et al*. (2022). Genes were considered ‘expressed’ in a layer if they showed coverage >1 transcripts per million (TPM) in at least 5 or more biological replicates. A gene was considered ‘induced’ if its median expression in the focal layer was at least 2x greater than the sum of its median expression in other two layers. Genes that were not induced in the junctional zone or labyrinth but had median expression in junctional and labyrinth greater than 2x median expression in the decidua were considered induced in the fetal placenta.

To assess whether genes were enriched for particular biological functions, we performed gene ontology (GO) enrichment analyses with the *gost* function of the R package gprofiler2 v 0.2.1 (Raudvere et al. 2019; Kolberg et al. 2020). This package uses hypergeometric tests for enrichment of categories from GO, KEGG, REACTOME, TF, MIRNA, and other databases within the subset of query genes, from which we excluded un-curated GO inferred from electronic annotations. We used the genes expressed in placental tissue as the background gene set and a Bonferroni correction for multiple testing.

### Detection of parent of origin effects in gene expression

We used allelic expression matrices to identify imprinted expression. First, we filtered expression matrices to identify genes suited for allele specific expression analysis on each cross type keeping genes with sufficient coverage (>10 reads) in at least four biological replicates overlapping at least two diagnostic variants separated by at least 150 bp in a reference transcript. Next, we tested for parent-of-origin expression bias at each gene using a *χ*^2^ test on the contingency table composed of the TPM normalized read counts of each allele in each direction of the cross at a p-value cutoff of 0.01 after Bonferroni correction. Additionally, we quantified the extent of parent-of-origin bias using the bias score proposed by Wang *et al*. (2011) as follows. First, the variables P1 and P2 were defined as the proportion of gene expression originating from a reference allele when inherited from the mother (P1), and the proportion of gene expression of the same allele when inherited from the father (P2) (*e*.*g*., in a dom x dom cross, P1 = % WSB allele expression in a ♀dom^WSB^ x ♂dom^LEW^ cross, and P2 = % WSB allele expression in a ♀dom^LEW^ x ♂dom^WSB^ cross). Bias scores were calculated as the difference between P1 and P2, which can range from -1 to 1 reflecting complete paternal and maternal bias respectively. A gene was considered biased if its absolute bias score was equal or greater than 0.4 corresponding to 70% or more expression originating from the maternally or the paternally inherited allele. This cutoff is the same used by Brekke et. al. Brekke *et al*. (2016) and slightly more conservative to other studies assessing parent-of origin expression bias in mouse placentas (65%, Wang *et al*. 2011; Andergassen *et al*. 2017).

### Estimation and correction of maternal contamination

Maternal contamination can be detected from deviations from 0.5 in the genome-wide proportion of RNA-seq reads assigned to the maternal allele among reads mapped to autosomal genes (Finn *et al*. 2014; Brekke *et al*. 2016). However, the contribution of maternal contamination to estimates of maternal expression in the placenta is not expected to be constant across genes, but proportional to a gene’s expression level in contaminating cells (Proudhon and Bourc’his 2010; Finn *et al*. 2014). Therefore, we developed a model to estimate and analytically correct for maternal contamination at individual loci. Unlike previous studies (Finn *et al*. 2014), our model does not rely on a subset of known paternally-expressed genes to measure contamination, and incorporates replicated measures of decidual expression levels specific to each maternal genotype.

Our strategy accounted for maternal contamination in two steps. First, we estimated the extent of maternal contamination using diagnostic gene-sets and samples (*i*.*e*., fetal placenta samples for which we kept decidual tissue) and next, we accounted for the effect of maternal contamination in each gene in all samples using estimates of median contamination and median gene expression levels in the decidua. To estimate the extent of contamination, we first modeled total gene expression in fetal placenta samples (*exp*_*Total*_) as the sum of gene expression from the maternally inherited allele (*exp*_*Mat*_), from the paternally inherited allele (*exp*_*Pat*_), and from maternal contamination (*exp*_*Cont*_)(equation 1). We next used a series of filtering criteria designed to identify a set of diagnostic genes with no allelic imbalance whose shifts from a 1:1 allelic expression ratio were most likely due to maternal contamination (see extended methods for details) and used this gene set to solve for *exp*_*Cont*_ in our model, assuming equal expression from the maternally and paternally inherited alleles (equations 2 and 3).

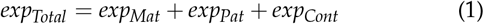

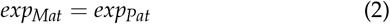

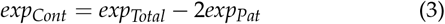

Estimates of *exp*_*Cont*_ from diagnostic gene sets (N varied by cross type; dom^LEW^ x dom^WSB^: 180, mus^PWK^ x mus^CZII^: 169, spret^STF^ x spret^SFM^: 201, mus^PWK^ x dom^WSB^:308) were then used to calculate median and maximum levels of maternal contamination (*C*), measured as the proportion of a gene’s expression level (TPM) in the decidua (*exp*_*Dec*_) required to explain contamination-derived maternal expression in the corresponding fetal placenta sample:

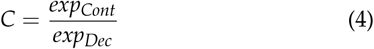

To account for the effect of maternal contamination across all expressed genes in all fetal placenta samples, we subtracted the theoretical contribution of maternal contamination (*exp* _*Dec*_ *\* C*) to the observed levels of maternal expression (*O*.*exp* _*Mat*_)in each gene using estimates of median *C* and median *exp*_*Dec*_ specific to each maternal genotype, and recalcualted P1 and P2 values on each sample (equations 5 and 6).

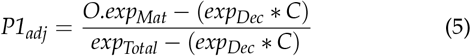

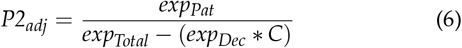

### Calling imprinted expression

After correcting for maternal contamination, genes with a *χ*^2^ corrected p-value below 0.01 and a parent-of-origin bias score greater than 0.4 or lower than -0.4 were considered maternally- or paternally-biased respectively (Wang *et al*. 2011; Brekke *et al*. 2016). To furhter reduce the number false positives, novel candidates for maternally-biased expression where only considered if they were in proximity (< 2Kbp) of a parent-of-origin differentially methylated region (see below), or if they met the following criteria: i) strong bias score (>0.7) under the most stringent threshold of contamination (maximum instead of median) in at least two cross types and ii) not primarily expressed in blood. The imprinted status of the genes in our final set of parent-of-origin biased genes was considered ‘confirmed’ for genes with known and validated imprinted status in the literature, or ‘candidate’ if the imprinted status of the gene has not been experimentally validated. Genes were considered ‘strong-candidates’ for imprinted expression if they were linked to a parent-of-origin DMR or if they were part of a known imprinted cluster. We refer to our final set (confirmed and candidates) as our ‘putatively imprinted gene set’, noting that candidates require further validation to be considered truly imprinted.

A reference list of genes with known and experimentally validated imprinted status was constructed by combining information from the Otago Imprinted Gene Catalog (https://corpapp.otago.ac.nz/gene-catalogue) and the Geneimprint databases (https://www.geneimprint.com). The list was manually curated to include updated records from recent publications (Andergassen *et al*. 2021, 2017; Richard Albert *et al*. 2023). Imprinted status of a gene was considered ‘validated’ if they had experimental support such as confirmation of imprinted expression from embryo transplant experiments, trophoblast cell cultures, or experimental manipulation of epigenetic marks associated with allele specific silencing.

### Estimation of divergence in parent-of-origin biased expression

To study divergence of imprinted expression, we performed all pairwise comparisons between the three *Mus* lineages using the subset putatively imprinted genes with sufficient power for allele specific expression in both lineages being compared. To detect changes in imprinted status, we looked for differences in the strength of parent-of-origin expression between lineages. We considered a gene divergent if the difference in score bias was greater than 0.4 between lineages.

### DNA-methylation analysis

DNA methylation analysis was performed using the *Methpipe* pipeline (Song *et al*. 2013). Filtered reads were mapped to the reference genome using *abysmal* (de Sena Brandine and Smith 2021), PCR duplicates were removed using the duplicate-remover script, and methylation levels were estimated at individual CpG sites using the *methcounts* and *symmetric-cpgs* scripts. Based on the resulting methylation calls, hypomethylated regions were identified using the *hmr* script and used to test for differential methylation using the *dmr* script with default parameters.

For allele specific methylation analyses, reads were first separated depending on their parent of origin. A custom python pipeline, *QuePadre*, was developed to identify reads of maternal or paternal origin based on informative polymorphic sites accounting for bisulphite transformation. To identify parent-of-origin differentially methylated regions, all CpG methylation calls originating from maternal reads and all CpG methylation calls originating from paternal reads across samples of a reciprocal cross were merged into compiled maternal and paternal CpG methylation calls using the *merge-methcounts* script (*e*.*g*., CpG methylation calls from PWK reads from all ♀mus^PWK^ x ♂mus_CZII_ samples and CpG methylation calls from CZII reads from all ♀mus^CZII^ x ♂mus ^PWK^ samples were merged to generate compiled maternal methylation calls). The resulting maternal and paternal methylation levels and hypomethylated regions were used for differential methylation analysis.

### Differential gene expression and network analysis

Differential gene expression analyses were performed with the R package *DESeq2* (Love *et al*. 2014). For every pairwise comparison, genes were kept for analysis if they showed expression levels of at least one TPM in at least three biological replicates. Genes were considered differentially expressed if they showed an absolute value of *log2FoldChange* (|*log2FC*|) of at least two and a maximum 0.01 adjusted p-value. To study overall patterns of placental gene expression, we constructed weighted gene correlation networks with the data from *M. m. musculus, M. m domesticus*, and their reciprocal hybrids using the R package WGCNA (Langfelder and Horvath 2008). Briefly, a soft threshold power of 24 was chosen based on network connectivity and used to automatically determine signed modules via dynamic cutting, with a merging threshold of 0.25. This process results in sets of genes that share similar expression patterns, placing up- and down-regulated genes into separate groups.

## Results

### Patterns of gene expression across placental layers reflect functional specialization at the transcriptome level

To gain insights into the evolution of layer specific transcriptomes, we identified and compared subsets of genes with induced expression in distinct placental layers from *M. m domesticus* and *M. spretus*. After filtering low quality RNA-seq reads, an average of 14.8 million RNA-seq reads per sample were mapped to annotated genes. In *M. m. domesticus*, 15,902 genes were expressed (TPM > 1) in the whole placenta, of which 4,259 (26.7%) showed induced expression. In *M. spretus*, 15,871 genes were expressed of which 3,138 (19.7%) showed induced expression. In both species, the decidua showed the greatest number of induced genes, followed by the labyrinth, and then the junctional zone (Figure 2A), Table S1).

**Figure 2.**
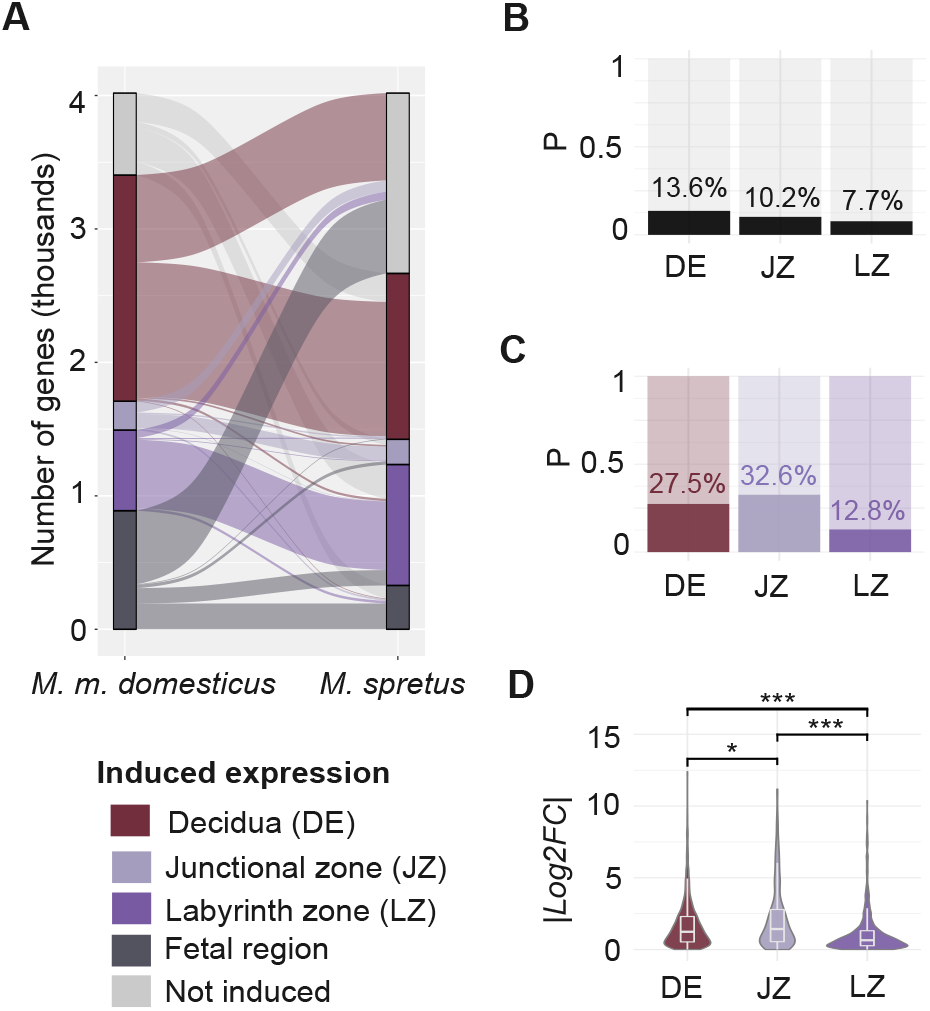
Interspecific divergence of layer-induced expression in the placenta. **A)** Alluvial plot showing overlap in sets of genes showing induced expression across placenta layers in *M. m. domesticus* and *M. spretus*. **B)** Barplot of the proportion of genes showing differential expression between species in the entire transcriptome of each placental layer. **C)** Barplots of the proportion of layer-induced genes showing differential expression between species **D**) Violin plots showing the distribution of |*log2FC*| scores across sets of genes with induced expression. Stars mark significance for a two-sided Wilcoxon ranked sum test in all pairwise comparisons.

In *M. m. domesticus*, we found enrichment of terms consistent with known layer specific functions among gene sets of induced expression. For example, 27 of the 40 enriched terms (67.5%) among decidua-induced genes were immune related with ‘immunoregulatory interactions between lymphoid and non-lymphoid cells’ (p-value = 5.91e-20), ‘cytokine-cytokine receptor interactions’ (p-value = 2.55e-16), and ‘complement and coagulation cascades’(p-value = 1.46e-07) among the top significantly enriched terms. In the junctional zone, seven of the eight enriched terms (87.5%) were related to signal transduction, including ‘class A/1 (Rhodopsin-like) receptors’ (p-value = 5.12e-10) and ‘neuroactive ligand-receptor interactions’ (p-value = 4.77e-07). Lastly, 11 of the 13 terms enriched among labyrinth-zone-induced genes (84.6%) were associated with nutrient transport, including ‘vitamin digestion and absorption’ (p-value = 3.25e-03), ‘transport of small molecules’ (p-value = 3.36e-03), and ‘regulation of insulin-like growth factor transport and uptake (p-value = 1.40e-02). A complete list of significant terms enriched on each gene set is provided in supplementary Table 2. Overall, these results demonstrate that the functional compartmentalization of the placenta is reflected in patterns of induced expression across layers, consistent with the evolution of layer specific regulatory programs that underlie tissue specialization.

Next, we compared evolutionary divergence of layer-specific transcriptomes between *M. m. domesticus* and *M. spretus*. Induced expression was partially conserved between species with nearly half (48%) of the genes induced in *M. m. domesticus* showing induced expression in the same layer in *M. spretus* (Figure 2A). However, this value varied by layer type with the highest similarity in the labyrinth zone where 513 of 738 genes (69.5%) showed the same pattern of induced expression between species, followed by the decidua (1,007 of 2,215 genes, 45.5%) and the junctional zone (105 of 319 genes, 32.9%). Differences in sets of induced expression between species were largely explained by gains and losses of expression or expression specificity but not by turnover of induced expression between tissues. For example, 103 of the 319 (32.2%) genes showing induced expression in the junctional zone of *M. m. domesticus* were not expressed in the placenta of *M. spretus*, 85 (26.6%) were not induced in any layer, and only 13 (4%) showed induced expression in other layers.

Comparisons of differential gene expression between species also revealed greater conservation in the labyrinth zone compared to the other layers when considering all expressed genes on each layer (Figure 2B) or only induced genes (Figure 2C). For induced genes, the junctional zone showed a greater proportion of differential expression (32.6%), followed by the decidua (27.6%) and the labyrinth zone (12.8%) (Figure 2C). The extent of expression level divergence among induced genes was also greater in the junctional zone (median |*log2FC*| = 1.43), followed by the decidua (median |*log2FC*| = 1.23), whereas expression levels were largely conserved among genes induced in the labyrinth zone (median |*log2FC*| = 0.64). Overall, these results suggest that interspecific divergence of placental gene expression levels and expression specificity is largely driven by regulatory divergence at the maternal-fetal interface in the decidua and the junctional zone.

### Maternally-biased genes predominate in the placenta after correcting for contamination

Our initial screen for imprinted expression in *M. musculus* detected significant maternal expression bias for 341 genes and paternal expression bias for only 18 genes using a moderate bias threshold of 0.4 (Wang *et al*. 2011; Brekke *et al*. 2016). Of these, 42 had validated imprinted status in mouse placenta, 124 had been previously detected in other RNA-seq based screens but lack experimental validation, and 199 were unique to this study. However, after model-based correction for maternal contamination, our final set consisted of only 49 maternally-biased genes along with the same 18 paternally-biased genes. This putative imprinted gene set include all 42 validated genes initially detected and 25 candidates that await validation, nine of which were unique to this study.

To evaluate the efficiency of our correction method, we searched for correlations between gene expression levels in the decidua and estimates of maternally biased expression in the fetal placenta, expected from contamination. If our mehtod effectively corrects for contaminaiton we expected these corelations to desappear after correction. Several observations supported this prediction. Our initial screen for imprinted expression showed a significant transcriptome-wide allelic skew towards maternal expression at autosomal loci (Exact Binomial, probability = 0.538 +/-0.004, p-value = 2.2e-16; Figure S4B), a positive correlation between decidual to fetal placenta expression ratios (D:F) and proportions of maternal allele expression (*pMat*) (Pearson’s correlation, r = 0.31, p-value = 2.2e-16; Figure S2), and a substantially larger skew towards maternal expression in the junctional zone compared to the labyrinth at loci with high decidual expression (junctional zone *pMat* = 0.69, labyrinth *pMat* = 0.52; Figure S4A). After correction, only minimal skew towards maternal expression remained (Exact Binomial, probability = 0.503 +/-0.001, p-value = 2.2 e-16) and the correlation between D:F and *pMat* in fetal placenta samples disappeared (Pearson’s correlation, r = 0.008, p-value = 0.38). Corrected bias scores also allowed us to recover the imprinting signal from the known paternally-biased gene *Impact*, originally lost in one of our crosses (mus^PWK^ x mus^CZII^) most likely due to the effect of maternal contamination. While not all false positives induced by contamination can be identified computationally due to potential violations of the assumptions of our simple model (*e*.*g*, equal expression levels in contaminating and non-contaminating decidual cells. See Discussion for details), overall, these results show that our model can help account for the effect of maternal contamination yielding a more robust representation of the placental imprintome.

Under the stringent cutoffs of contamination of our correction method, maternally-biased genes comprise 73% (49 genes) of our final set, supporting a prevalence of maternally-biased expression in the placenta. We were unable to confirm imprinted expression of 11 genes with validated imprinted status in the mouse placenta literature, of which three were expressed at low levels (*Th, Magi2, Rasgrf1*), four were expressed but not biased (*Sall1, Cd81, Park2, Smoc1*) and five showed bias below the 0.4 bias score threshold (*Axl, Phactr2, Phf17, Arid1b, H13*).

### Large contributions of maternal DNA-methylation to parent-of-origin expression in the placenta

We next compared patterns of DNA methylation of the maternal and paternal genomes in fetal placenta samples from two reciprocal *M. musculus* crosses (mus^PWK^ x mus^CZII^, mus^PWK^ x dom^WSB^). We used these data to discover new putative control regions of imprinted expression and measure the contribution of maternal and paternal methylomes to the regulation of parent-of-origin expression in the placenta. A total of 3,947 differentially methylated regions (DMRs) were identified, of which, 602 were linked (within 2 Kbp of gene boundaries) to genes expressed in the placenta. Our analysis recovered overlap between parent-of-origin DMRs and known imprinted loci in the mouse genome (Wang *et al*. 2014), with 5 parent-of-origin DMRs link to a total of 22 known imprinted loci including 16 known imprinting control regions (Table S). We then looked for support of genomic imprinting at the 25 candidates detected through our screen of allele-specific transcript bias. We identified five DMRs linked to candidates for imprinted expression (*Mamdc2, Adamts5, Slco5a1, Sez6l, Tdo2*). We therefore considered these 5 genes strong candidates for imprinted expression.

A vast majority of DMRs (95%) were determined by maternal DNA methylation (Figure 3A) and, although most maternal DMRs were not linked to expressed genes, maternal DNA methylation was also prevalent among imprinted loci. Of 4,740 maternally methylated DMRs, 499 (13.3%) were linked to genes expressed in the placenta of which 23 showed significant parent-of-origin expression bias (Figures 3B,C). In contrast, of 207 paternally methylated DMRs, 103 (49.7%) were linked to expressed genes, but only three of those showed significant parent-of-origin expression bias (Figure 3B). Notably, maternally-methylated DMRs were linked to both maternally- and paternally-biased expression, including 13 linked to known paternally-biased genes, five linked to known maternally-biased genes and 5 linked to novel candidates for maternally-biased expression (Figure 3C). Regulatory mechanisms underlying simultaneous DNA-methylation and increased expression of maternal alleles at these candidate loci was not clear from DNA methylation patterns as most maternally methylated regions did not overlap annotated regulatory regions in the mouse ENCODE project (Figure S5).

**Figure 3.**
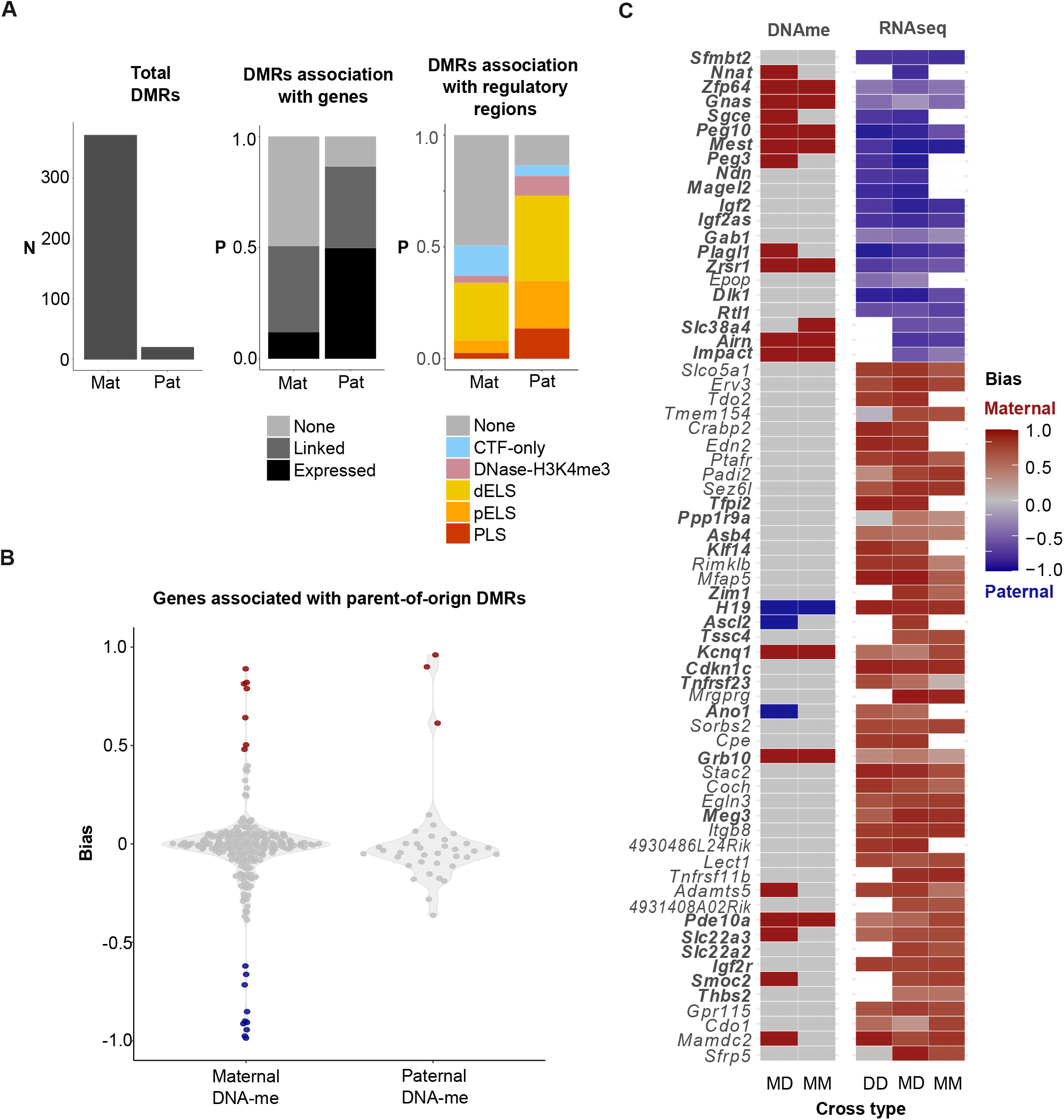
Maternally methylated regions predominate among imprinted loci. **A)** Functional characterization of DMRs of maternal and paternal origin in the mus x dom cross. **B)** Distribution of bias scores of genes associated with maternal and paternal DMRs in the mus x dom cross. **C)** Overlap between parent-of-origin DMRs and genes showing parent-of-origin expression bias in RNA-seq datasets. Each column in the tile plot is labeled according to the cross type as follows: MM: mus x mus, DD: dom x dom, MD: mus x dom

### Layer-specific enrichment of genes with parent of origin expression

Having defined a core set of imprinted genes, we asked whether tissue specialization influences the evolution of imprinted expression in the mouse placenta. We found that 31 genes from our putatively imprinted set (43%) were significantly overrepresented among the 1,057 genes showing induced expression in fetal placental layers (Fisher’s Exact, odds ratio: 9.27, p-value = 2.2e-16). Next, we tested for enrichment of imprinted expression among genes with induced expression in the junctional and labyrinth zones separately. We found a strong pattern of layer-specific enrichment of imprinted genes, with maternally-biased genes preferentially expressed in the junctional zone, (15 genes, Fisher’s Exact, odds ratio = 17.8, p-value = 5.3e-13) and paternally-biased genes preferentially expressed in the labyrinth zone (9 genes, Fisher’s Exact, odds ratio = 12.01, p-value = 1.14e-06; Figure 4). This striking polarization of parent-of-origin expression bias in the placenta suggests that parent-of-origin effects on gene expression are not randomly distributed across placental layers and that distinct placental functions may influence the evolution of a maternally- and paternally-biased expression.

**Figure 4.**
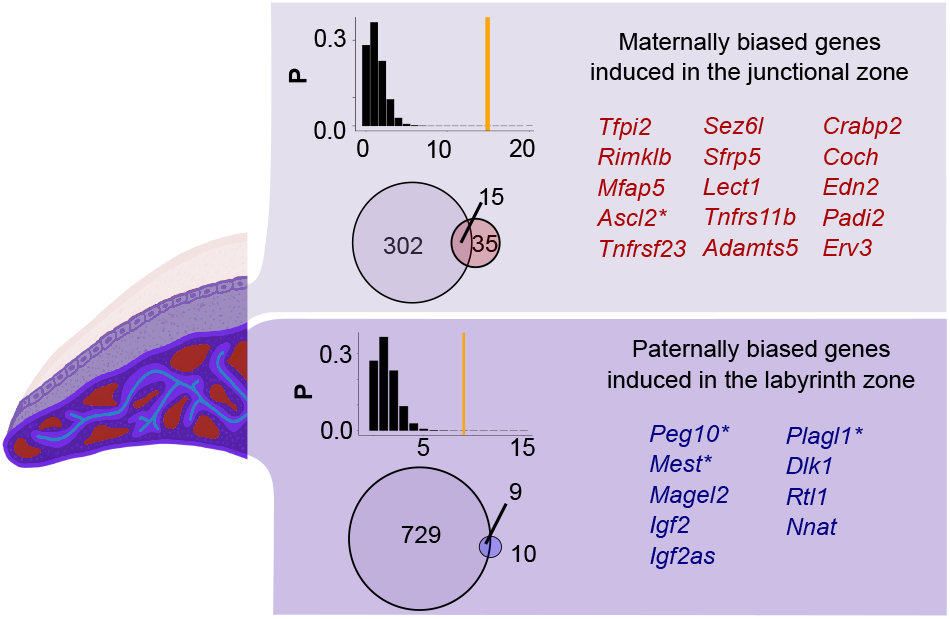
A subset of imprinted genes are associated with placenta tissue specialization. On the left, a schematic representation of placental layers. On the right, the box on top shows the overlap between genes showing induced expression in the junctional zone and maternally-biased expression. The Venn diagram shows the overlap between the two gene sets, and the bar plot shows the probability of the observed overlap size estimated from the hypergeometric distribution. Box at the bottom shows overlap between genes showing induced expression in the labyrinth zone and paternally-biased genes. Genes in bold have validated imprinted status in the literature, and genes marked with an asterisk are genes with imprinted status supported by differential methylation between maternal and paternal genomes in this study.

### Extensive co-regulation of imprinted genes in the placenta

After identifying parent-of-origin and induced expression in the placenta, we were interested in exploring potential regulatory links among genes of interest. To that end, we performed a weighted gene correlation network analysis (WGCNA) to cluster genes based on co-variation of expression levels across fetal placenta samples from *M. m. musculus, M. m. domesticus*, and their reciprocal crosses (59 samples passing filter) and tested for enrichment of gene-sets of interest across the resulting gene modules.

After filtering lowly expressed genes, we analyzed 15,863 genes expressed (TPM > 1) in the fetal placenta and identified 22 modules of co-expression, ranging in size from 43 to 2,622 genes. Two of these modules stood out for their enrichment of imprinted expression after correcting for maternal contamination. The first of these (Figure 6, module “E”), was a small module (43 genes) enriched for maternally-biased genes (four genes, Fisher’s Exact p-value = 0.003) and junctional-zone-induced genes (11 genes, Fisher’s Exact p-value = 5.9e-10). Only six terms were represented in this module (‘homeostatic process’, ‘cell differentiation’, ‘anatomical structure development’, ‘signal transduction and mitotic cell cycle’) but none were overrepresented at a p < 0.05 threshold.

The second module (Figure 5, module “G”, 297 genes) was also enriched for maternally-biased genes (14 genes, Fisher’s Exact p-value = 9.17e-11) and junctional-zone-induced genes (18 genes, Fisher’s Exact p-value = 9.5e-6), as well as for decidua-induced genes (79 genes, Fisher’s Exact p-value = 2.3e-9) and X-linked genes (23 genes, Fisher’s Exact p-value = 4.7e-4). All GO terms enriched in this module were associated to immunological responses such as antigen processing and presentation of endogenous peptide antigen (p-value = 1.5e-4), regulation of cell killing (p-value = 1.5e-4), and regulation of T cell (p-value = 1.58-4), leukocyte (GO:0002703, p-value = 2.25e-4) and lymphocyte (GO:0002706, p-value = 2.25e-4) mediated immunity. Thus, this module appears to reflect a regulatory network associated with the modulation of the maternal immune system.

**Figure 5.**
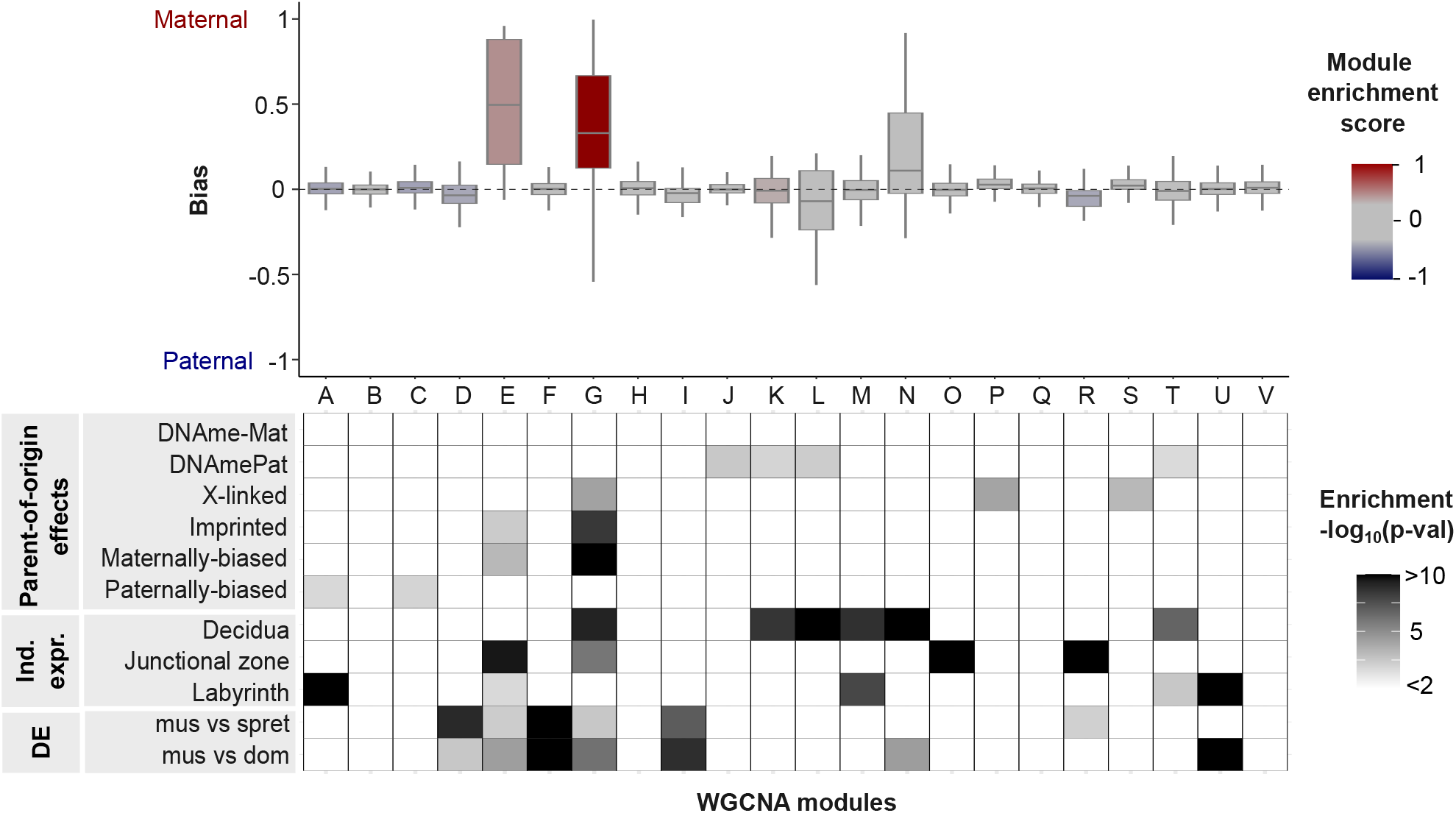
Enrichment of target gene sets across WGCNA modules. 22 gene modules identified through WGCNA show differences in the distribution of parent-of-origin expression bias and enrichment of imprinted and induced genes. Boxplot at the top shows distributions of parent-of-origin bias scores among genes on each module. Box colors reflect enrichment for imprinted expression using our putative imprinted gene set as a reference. Imprinted enrichment score ranges from 10 to -10 and is calculated as the -Log10 of Fisher’s exact p-value multiplied by -1 if the bias is towards paternal expression. Tile plot at the bottom shows enrichment of all gene sets of interest genes with parent-of-origin effects on gene expression and DNA methylation, layer-induced genes and differentially expressed genes.

We noticed that besides enrichment for parent-of-origin and tissue biased genes, the “G” module was also enriched for genes showing differential expression, but appeared dominated by genes with differences in expression levels between *M. m. musculus* and *M. m. domesticus*. Therefore, we further explored the subset of DE genes contained in this module (63 genes) to gain insights into the potential roles of tissue specialization and parent-of-origin expression in evolutionary divergence of placental tissues. Of 63 DE genes in this group, 34 showed a derived expression level in *M. m. domesticus*, but none of these were part of our putative set of imprinted genes. Instead, divergence at these loci appeared to be tightly linked to regulatory divergence in the decidua: 12 of these genes showed induced expression in the decidua and all 34 genes showed strong downregulation specific to *M. m. domesticus* and accentuated in the decidua (Fig. S6). This pattern suggests rapid divergence of the placental transcriptome in *M. m. domesticus* associated with the evolution of a maternal regulatory network.

Lastly, we were interested in evaluating whether our method for addressing maternal contamination allowed us to identify sets of coexpressed genes that were particularly susceptible to maternal contamination. Encouragingly, 17 of the 22 modules showed an adjusted median bias score centered at or near zero after correction, indicating that our model effectively corrected these gene sets for maternal expression. Two modules (“L”,and “R”) became slightly paternally-biased, which may indicate that our model is overcorrecting for maternal contamination in these genes. Of note was a set of genes that was heavily biased towards maternal expression, but centered near zero after correction (161 genes, “N” module, Fig. S7). This module showed high intramodular connectivity (Fig. S8), indicative of a high degree of covariation among genes. Importantly, most genes in this module (112, 70%) showed induced expression in the decidua, thus, signals of covariation in the fetal placenta most likely reflect coordinated expression in maternal cells. Consistent with this, GO enriched terms for this module all suggest maternal immune function, with the macrophage scavenger receptor 1 as the most connected gene in the network. This suggests that these genes are not a random assortment of transcripts scattered in residual maternal tissue, but instead may belong to a gene regulatory network from maternal immune cells. Interestingly, this module showed significant differences in the extent of maternal bias between lineages (Fig S9 ANOVA F(3,630) = 8.427, p < 0.0001, significantly lower in *M. m. musculus* by TukeyHSD) suggesting variation in the extent of maternal expression of this module between lineages.

### Rapid regulatory divergence at imprinted loci in the *M. m. domesticus* lineage

To gain insights into the extent of divergence at imprinted loci between closely related species, we performed pairwise comparisons of parent-of-origin expression bias and expression levels in our putative set of imprinted genes. Unexpectedly, divergence in parent-of-origin bias did not appear to scale with phylogenetic distance. Comparison between *M. m. musculus* and *M. m. domesticus* (31 genes) revealed differences above our threshold in parent-of-origin bias scores at three loci, including the known imprinted gene *Tnfrsf23* which showed significant maternal bias in *M. m. domesticus* and but not in *M. m. musculus* (Figure 6A). In contrast, our comparison between *M. m. musculus* and *M. spretus* (32 genes) did not reveal differences in imprinted expression (Figure 6B). To polarize the divergence observed between *M. musculus* lineages we also compared bias scores between *M. m. domesticus* and *M. spretus* (33 genes) which revealed three genes showing bias score differences above our threshold, including *Tnfrsf23* and *Rimkbl* which showed weak bias in *M. spretus*. These two genes also showed weak or no bias in *M. m. musculus*, suggesting their strong maternal bias is a derived state in *M. m. domesticus*.

**Figure 6.**
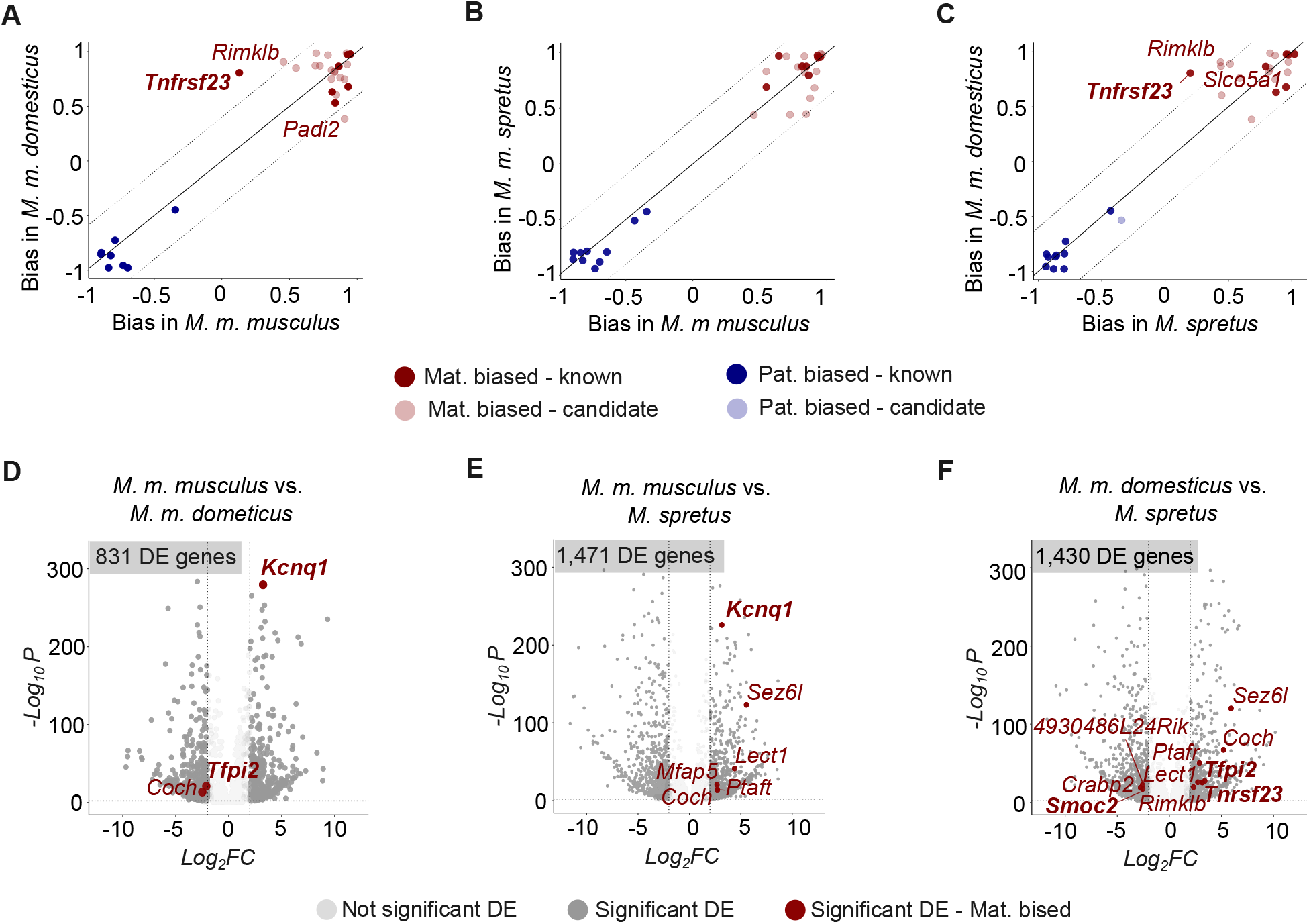
Rapid regulatory divergence of parent-of-origin biased genes. **A-C**: Pairwise comparisons of parent-of-origin bias scores of genes in our putatively imprinted set. Solid diagonal line follows the trajectory of equal bias in the two lineages. Dotted diagonal lines delimit a divergence threshold corresponding to score bias differences greater than 0.4 between lineages. Comparisons are limited to the subset of parent-of-origin biased genes with power for measuring allele specific expression in both lineages being compared. **D-F**: Volcano plots showing the results from pairwise differential expression tests highlighting genes with parent-of-origin expression. Box at the top left corner shows the number of significantly differentially expressed genes. Darker points correspond to known imprinted genes and lighter points to candidates

Differential expression tests also revealed greater expression level divergence at imprinted loci in *M. m. domesticus* than in *M. m. musculus* when compared to *M. spretus*. A total of 12 genes with parent-of-origin expression showed significant differences in expression levels in at least one pairwise comparison, of which, three were differentially expressed between *M. m. musculus* and *M. m. domesticus*, six between *M. m. musculus* and *M. spretus*, and 10 between *M. m. domesticus* and *M. spretus*. Of note, the *Kcnq1* imprinted cluster showed evidence of repeated instances of regulatory divergence. Within this cluster, the *Kcnq1* gene was downregulated in *M. m. musculus* compared to the two other lineages, and *Tnfrsf23*, which showed a derived imprinted status in *M. m. domesticus* (see above), had lower levels of expression in *M. m. domesticus* when compared to *M. spretus*. Also worth noting was the abundance of junctional-zone-induced genes among parent-of-origin genes showing differential expression (7/12, 58%) suggesting a role of junctional zone specialization in the evolutionary divergence at parent-of-origin loci in the placenta.

## Discussion

We generated genome-wide gene expression and DNA-methylation data from three closely related lineages of house mice to study the evolution of parent-of-origin expression across distinct functional layers of the mouse placenta. This comparative dataset allowed us to uncover important links between core elements of placental biology and the evolution of genomic imprinting. We detected a strong polarization of parent-of-origin effects across placental layers with enrichment of paternally-biased expression in the labyrinth zone and enrichment of maternally-biased expression in the junctional zone. We also identified a putative regulatory network enriched for maternally-biased expression that is likely involved in immunomodulation and shows rapid divergence in the *M. m. domesticus* lineage. These results provide novel insights into the role of tissue specialization and maternal-fetal interactions in the evolution of imprinted expression in the mouse placenta.

### Detection of imprinted expression, maternally-biased expression, and contamination

Genome-wide screens of allele specific expression have been instrumental for identifying imprinted expression in mammals (*e*.*g*., Wang *et al*. 2011, 2013; Babak *et al*. 2015; Brekke *et al*. 2016), but, are also notoriously susceptible to experimental and bioinformatic errors (Proudhon and Bourc’his 2010; DeVeale *et al*. 2012). While many of these issues apply generally across tissue types (*e*.*g*., noise associated detection of allele-specific expression), the intimate interactions between fetal and maternal tissue presents unique challenges to the detection of imprinting in the placenta. Even moderate amounts of remnant maternal cells in fetal placental samples can inflate estimates of maternally-biased expression (Proudhon and Bourc’his 2010; Okae *et al*. 2012), sparking skepticism over the commonly observed excess of maternally-bias genes in the placenta (Proudhon and Bourc’his 2010; Wang *et al*. 2011; Okae *et al*. 2012). Nonetheless, of all mammalian tissues, the placenta is arguably the most important for understanding the evolution of genomic imprinting given the close association between the evolution of placentation and imprinting (Killian *et al*. 2000, 2001; Edwards *et al*. 2008), its susceptibility to genetic conflict (Moore and Haig 1991), and the critical roles that imprinted genes play in early development (Thamban *et al*. 2020).

The relative abundance of maternally-biased genes in the placenta has varied by experimental strategy and taxonomic group. For example, studies that have modeled decidual contamination in the mouse placenta (Finn *et al*. 2014), as well as screens in other rodent systems with less invasive placentation (Brekke *et al*. 2016), have identified a strong excess of maternally-biased genes (> 90%). In contrast, screens at earlier stages of placental development (textiti.e., before fetal invasion; Andergassen *et al*. 2021), efforts to avoid maternal contamination through targeted placenta dissections (Wang *et al*. 2011) and screens from isolated trophoblast cells (Calabrese *et al*. 2015), all suggest more equal representation of maternally- and paternally-biased genes. Indeed, screens from species with non-invasive placentation such as equines found an excess of paternally-biased genes (Wang *et al*. 2013). These patterns could be interpreted as strong evidence that an excess of maternal bias in some studies likely results from contamination; however, they could also reflect evolutionary differences in patterns of imprinting between different mammalian lineages, across developmental time points and cell types, or both. Given the difficulties imposed by maternal contamination, RNA-seq based screens of imprinted expression are most powered when accompanied by independent signatures of allele specific DNA methylation or open chromatin that should be less sensitive to contamination.

Our study, combining elements of experimental and epigenomic validation with strain and species replication, supports an excess of maternally-biased expression in the rodent placenta. We found that ∼70% of genes with parent-of-origin expression were maternally-biased after correcting for contamination. Our estimates of maternal contamination were comparable to those from a similar study of the mouse placenta (Finn *et al*. 2014,; our range = 0.08% to 16% vs 0% to 12%). Our putative set of imprinted genes was robust across species and largely consistent with literature reports (Wang *et al*. 2011; Okae *et al*. 2012; Finn *et al*. 2014; Andergassen *et al*. 2017,;Figure S10), containing 42 known imprinted genes and 16 genes previously reported in mice. However, false positives for imprinted expression are expected to escape model-based correction if they violate the assumption that gene expression levels in contaminating cells can be accurately estimated from expression levels from bulk decidua tissue. For example, signals from the trophoblast tissue can disproportionately alter gene expression in the decidual cells closest to the maternal-fetal interface (Clark *et al*. 2002), which may lead to gene expression profiles in contaminating cells that differ from the bulk of the maternal decidua.

Our results are also consistent with previous studies concerned with maternal contamination and may help conciliate conflicting results in the literature. Our initial (uncorrected) screen detected maternal bias in 19 genes whose imprinted status was tested in the mouse placenta using embryo transplants and trophoblast cell cultures (Okae *et al*. 2012). Of these, eight were confirmed as imprinted and five as false positives in both studies, while seven were identified as false positives only in their study. This partial discrepancy could reflect the limitations of our model-based correction but also, experimental limitations such as the potential for disruption of epigenetic pathways induced by embryo transplant (*e*.*g*., Barberet *et al*. 2021). We also observe enrichment of paternally biased genes in the labyrinth zone which could explain the excess of paternally-biased genes observed in a previous study that experimentally avoided sampling tissue near the junctional zone (Wang *et al*. 2011). If paternal expression predominates in specific cell lineages from the labyrinth zone, dissections targeting the fetal side of the placenta may be enriched for labyrinth trophoblast cells, thus, enriched for paternally-biased expression. Single cell evaluation of imprinted expression in the placenta could help confirm this hypothesis.

Regardless of the specific mechanism responsible for maternal bias in unvalidated genes, we highlight the potential biological relevance of their expression patterns. Of 27 imprinted candidates showing maternal bias, over half (15) were clustered in a WGCNA module with known imprinted genes, X-linked genes, and genes with induced expression in the maternal decidua. Notably, this putative regulatory network was also enriched for immune functions associated with antigen presentation. Additionally, five of the 15 imprinted candidates in this module (*Erv3, Edn2, Mgat3, Rnf182, Mfap5*) showed induced expression in the junctional zone and encode membrane or extracellular signaling proteins associated with trophoblast invasion, immunomodulation, and maternal-fetal communication (Marchand *et al*. 2011; Pavličev *et al*. 2017; Deng *et al*. 2019). These results support a role of maternally-biased expression in facilitating maternal-fetal interactions during development, as predicted by some theoretical models of placental regulatory evolution (Wolf and Hager 2006; Patten *et al*. 2016,; more below). Notably, these genes are robust to our stringent correction and their maternal bias is replicated across species and studies. Regardless of whether bias is the result of imprinting or expression from maternal cells functionally integrated with the junctional zone, maternally-biased expression at these loci is likely important for placental development, function, and evolution.

### Tissue specialization and the evolution of placental gene expression

As the site of nutrient transfer during development, the placenta is considered an arena for evolutionary conflict due to distinct fitness optima of maternally and paternally inherited alleles over offspring resource allocation (Moore and Haig 1991; Haig 2000). However, the mammalian placenta performs several functions beyond the regulation of nutrient transfer, including modulation of the maternal immune response, regulation of adaptive changes in maternal metabolism, and efficient molecular cross-talk of maternal and embryonic tissues for successful implantation and development (Carter 2012). It is unclear, however, how distinct functions have influenced the evolution of the mammalian placenta at the molecular level.

Our results provide new insights into the roles of tissue specialization and functional compartmentalization to the evolution of gene expression and genomic imprinting in the mouse placenta. Our layer enriched expression analyses revealed functional specialization at the transcriptome level, with 4,259 and 3,138 genes showing layer induced expression and GO enrichment for layer-specific functions in *M. m. domesticus* and *M. spretus*, respectively. In parallel, expression divergence between *M. m. domesticus* and *M. spretus* revealed two important patterns in the evolution of tissue specialization. First, divergence in patterns of induced expression between species appears to occur through independent gains and losses of expression specificity in the placenta. Second, divergence in expression levels and specificity was elevated at the decidual-fetal placenta interface compared to the labyrinth zone. This complements findings of other studies that have identified evidence of rapid molecular divergence in the junctional zone in rodent placentas. For example, a study on protein sequence evolution in rodent placentas found that the most rapidly evolving placental proteins were hormones secreted by the fetal placenta that directly influence maternal physiology and maternal immune response (Chuong *et al*. 2010). Additionally, most differentially methylated DNA regions between mouse species are concentrated in the junctional zone (Decato *et al*. 2017). Together, these results provide evidence of rapid evolution of protein sequences, gene expression, and DNA-methylation in the junctional zone, suggesting that this layer plays an important role in the evolution of rodent placentas.

Tissue specialization also appears to be associated with patterns of imprinted expression in the mouse placenta. The strong polarization of parent-of-origin expression across placental layers suggests that distinct placental functions favor imprinting in different directions with prevalence of maternally-biased expression in the junctional zone and paternally-biased expression in the labyrinth zone. Genomic imprinting is commonly considered the result of evolutionary conflict between maternally and paternally inherited alleles over offspring resource allocation (Haig and Graham 1991). This evolutionary model, formally described as the kinship theory of genomic imprinting (Moore and Haig 1991; Haig 2000) predicts paternally-biased expression at loci that promote growth and maternally-biased expression loci that restrict growth. Consistent with this, our study identified several growth enhancers among paternally-biased genes as well maternal DNA epigenetic modifications known to be involved in the silencing of the maternal allele at these loci. Most paternally-biased genes showing labyrinth-zone-induced expression (seven of nine) are known to promote cell proliferation and angiogenesis in the placenta (*Peg10, Mest, Igf2, Dlk1, Rtl1, Plagl1, Nnat*; Mayer *et al*. 2000; Constância *et al*. 2002; Sekita *et al*. 2008; Tunster *et al*. 2018; Xie *et al*. 2018; Starks *et al*. 2020; Xing *et al*. 2022). This could reflect a mechanism to limit resource allocation through maternal imprints that restrict angiogenesis and cell proliferation at the main region of nutrient transfer in the placenta. In contrast, maternally-biased genes showed a more diverse functional landscape including growth restriction, immunomodulation, and cell-cell adhesion, and were overrepresented among junctional-zone-induced genes. Of the 15 genes showing maternal bias and induced expression in the junctional zone, five are known to function as tumor suppressors (*Sez6l, Tfpi2, Ascl2, Lect1, Adamts5*) (Hiraki *et al*. 1999; Kumar *et al*. 2012; Tunster *et al*. 2016; Xiao *et al*. 2017). Allelic silencing of genes that inhibit cell proliferation in the junctional zone may serve as a mechanism to promote embryonic growth by facilitating proliferation of glycogen storage trophoblast cells that fuel late development (Esquiliano *et al*. 2009), also consistent with predictions of the conflict theory.

We also observed several other functional, epigenetic, and gene expression patterns that may be more clearly predicted by alternative models of imprinted expression evolution suggesting that placental functions other than nutrient transfer may contribute to the evolution of imprinted expression in the junctional zone. The mother offspring coadaptation theory (Wolf and Hager 2006) conceives genomic imprinting as a mechanism that facilitates developmental integration of the maternal and fetal genomes by reducing negative epistatic interactions between the maternal and paternal gene copies in the placenta. Consistent with this, we observed prevalent imprinted expression in the same direction among interacting genes (Patten *et al*. 2016) and maternally-biased expression at loci that are directly involved in intimate interactions between maternal and fetal tissues (Wolf and Hager 2006). One of the most striking patterns from our WGCNA analysis was that the main gene set enriched for maternally-biased and junctional-zone-induced expression was also enriched for X-linked and decidua-induced genes. In rodent placentas, X chromosome inactivation is not random but parent-of-origin dependent with the paternal X uniformly silenced across cells (Takagi and Sasaki 1975; Vrana *et al*. 2000; Brekke *et al*. 2016; Richard Albert *et al*. 2023). Thus, this putative regulatory network is broadly enriched for maternally-biased expression in the fetal placenta at loci predicted to interact with genes expressed from maternal cells, consistent with a role of maternally-biased expression in the integration of maternal and embryonic regulatory networks. A role for maternal-fetal communication in the evolution of placental imprinted expression was also supported by the functional composition of this gene set. Five of the 15 genes showing maternally-biased and junctional-zone-induced expression (*Erv3, Edn2, Tnfrsf23, Tnfrsf11*, and *Coch*) are known to be expressed in trophoblast cells and modulate maternal immune responses during development (Elferink and de Koster 1996; Clark *et al*. 2002; Py *et al*. 2013; Denner 2016). Additionally, the genes *Crabp2, Mfap5*, and *Ascl2* are known to mediate endometrial-trophoblast interactions during invasion (Enquobahrie *et al*. 2011; Zheng *et al*. 2022; Marchand *et al*. 2011). Remarkably, *Tnfrsf23* shows a unique expression pattern in the placenta with expression limited to a subpopulation of maternal and fetal cells that are in direct contact with each other in the placenta (Clark *et al*. 2002), suggesting a role in the modulation of molecular cross-talk between maternal and fetal tissues. Evidence of co-regulation among many of the same maternally-biased autosomal and X-linked genes has also been reported in the hamster placenta (Brekke *et al*. 2021), further supporting the biological relevance of maternally-biased expression in rodent development.

Lastly, we also observed overrepresentation of epigenetic modifications of the maternal genome among DMRs linked to imprinted loci, including 19 known maternally- and paternally-biased genes and five novel DMRs linked to maternally-biased genes. Overrepresentation of maternal DMRs has been observed in mice (Kobayashi *et al*. 2012), rats (Richard Albert *et al*. 2023), and humans (Hanna *et al*. 2016) but this excess of maternal DMRs appears limited to the placenta when compared to other organs (Hanna *et al*. 2016). This pattern may suggest more extensive control of maternal DNA methylation over imprinted expressison in extraembryonic tissues. In support of this, experimental deletions of a differentially methylated region in the *Dlk1-Gtl2* imprinted cluster in the mouse placenta showed loss of imprinted expression in both paternally- and maternally-biased genes of the cluster when the deletion was inherited from the mother, and no effects when the deletion was inherited from the father (Lin *et al*. 2003). These patterns are difficult to explain under an arms-race model of antagonistic coevolution (but not impossible, see Reik and Walter 2001; Haig 2014) and could reflect an epigenetic mechanism in support of the hypothesized extensive control of the maternal genome over imprinted expression in the placenta (Wolf and Hager 2006).

### Evolutionary divergence of the imprinted landscape

Regulatory divergence of imprinted genes in the placenta is believed to contribute disproportionately to the evolution of reproductive incompatibilities in mammals (Vrana 2007; Crespi and Nosil 2013). However, due to the lack of data on the placental imprinted landscape in most mammal species, it is unclear how rapidly imprinting evolves in nature.

Our comparative analysis of imprinted expression in three closely related mouse lineages provides new insights into the evolutionary divergence of placental imprinted expression between closely related species. Generally, we observed extensive conservation of imprint status at most genes evaluated, with turnover of imprinting limited to just one gene (*Tnfrsf23*) that showed biallelic expression in *M. m. domesticus*. Three other genes showed differences in the extent of bias (*Rimkbl, Padi2, Slco5a1*) but with no evidence for complete loss or gain of imprinted status in any comparison. A recent comparison between mouse and rat suggests that non-canonically imprinted genes contribute disproportionately to interspecific divergence in imprinting (Richard Albert *et al*. 2023). We did not detect a significant difference in parent-of-origin expression bias among any of the nine genes known to be non-canonically imprinted in mice. However, we had limited power to detect imprinted expression at these loci, as three genes did not show sufficient expression levels or had insufficient informative variants. Interestingly, of the five genes that had power for allele specific expression, we only confirmed expression bias in three (*Sfmb2, Gab1, Slc38a4*) and observed biallelic expression in two (*Sall1, Smoc1*). These genes were previously identified as imprinted at earlier developmental stages (Inoue *et al*. 2017b), and are known to rely on secondary methylation in the zygote to maintain their noncanonical imprints (Chen *et al*. 2019). Our lack of power due to low expression levels in mature placenta, and the observed biallelic expression in two of these genes, may reflect a transitory nature of non-canonical imprints limited to early developmental placental stages. We also observed several imprinted genes showing significant differences in expression levels among lineages. Among these, we highlight the divergence in expression levels of *Kcnq1* and *Tnfrsf23* genes, both located in the large *Kcnq1* imprinted cluster and recently reported to exhibit transgressive (outside of the range of expression of both parental lines) DNA-methylation and expression in *M. m. domesticus* x *M. spretus* hybrid placentas (Arévalo *et al*. 2021). Taken together, these results suggest that regulatory divergence in the *Kcnq1* imprinted cluster may contribute to the emergence of reproductive incompatibilities between these species.

In summary, our comparative analysis of parent-of-origin gene expression and tissue specialization revealed fundamental aspects of placental regulatory evolution. We have demonstrated how tissue specialization in the mouse placenta influences patterns of molecular evolution, including differences in rates of gene expression level divergence and patterns of imprinted expressions across functionally distinct layers. The evolution of imprinted expression in mammals is intricately associated with the evolution of unique aspects of placental biology, including the evolution of novel cell lineages and interactions between maternal and fetal cells (Renfree *et al*. 2009, 2013). Future studies on imprinted expression in the context of cell-specific biology can further illuminate fundamental aspects of the evolution of this important form of gene regulation in mammals.

## Data availability

All raw sequencing reads are archived at NCBI under BioProject XXXX. Accession numbers for individual libraries are provided in Supplementary Table SX. All custom scritps designed are avaialable in the Github repository for this project (https://github.com/LF-Rodriguez/Mouse_imprinting_evolution). Supplementary material is available at XX

## Acknowledgements

We would like to thank Kathryn Wilsterman and Polly Campbell for their contributions to the study design and interpretation of results. Zac Chevron, Lila Fishman, Brandon Cooper, Cynthia Ulbing, Nathaniel Herrera, Ana Paula Arprigio, and Jackie Olexa, provided helpful feedback throughout the development of the study. Kathryn Wilsterman, Polly Campbell and Erica Larson provided comments on an earlier version of this manuscript. Emily Kopania, Schuyler Liphardt and Gregg Thomas provided advice on the bioinformatic analyses. Sebastian Mortimer, Kelly Carrick, Pam Broussard and the University of Montana LAR staff helped with animal care.

