## Supplementary material for "Evolution of parent-of-origin effects on placental gene expression in house mice": Suplemental figures

2023

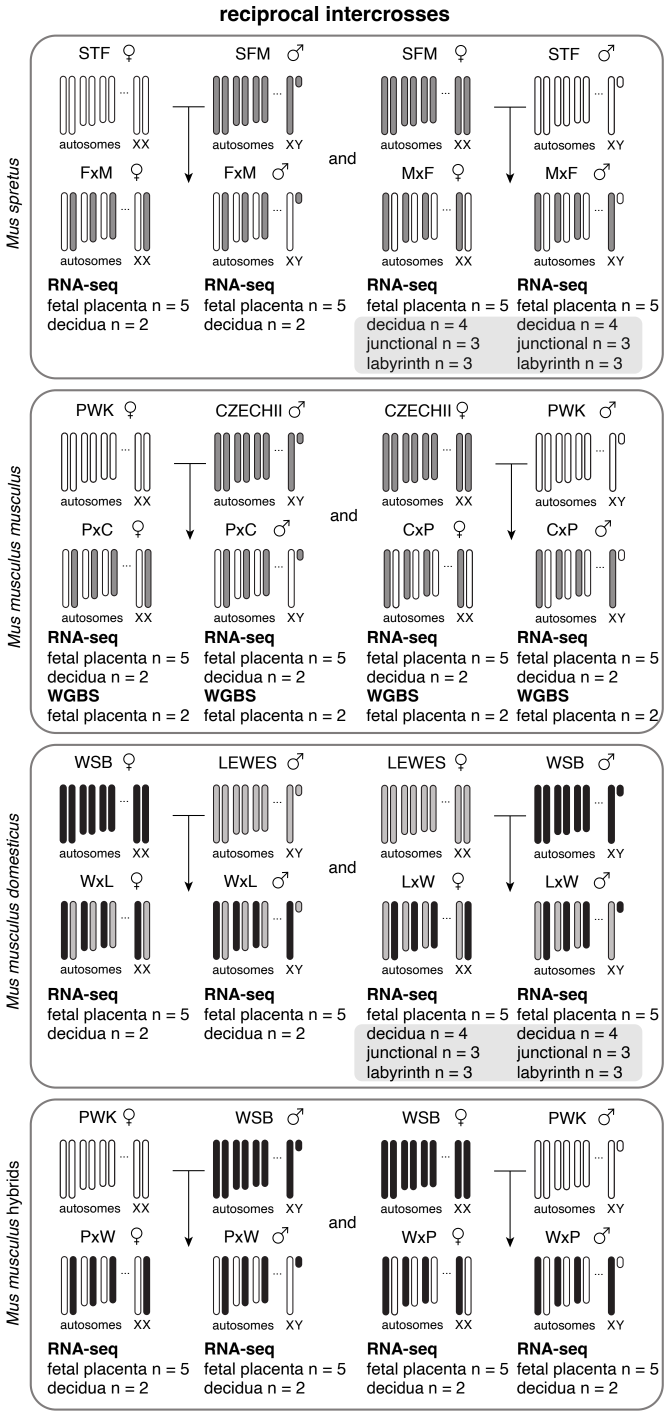


Figure 1: **Experimental crosses and sampling of genomic data**. Design of reciprocal crosses from three mus lineages used in this study, indicating specific wild derived inbred lines (PWK, CZECHII, WSB, LEWES, STF, and SFM) with sample sizes for each data type, mRNA sequencing (RNA-seq) and whole genome bisulfite sequencing (WGBS).

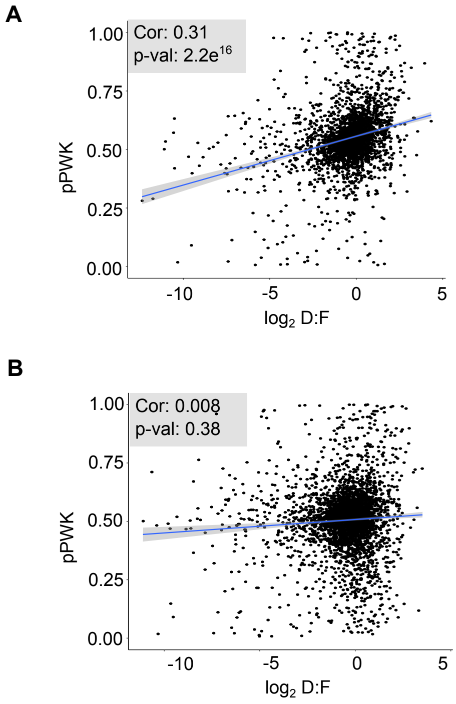


Figure 2: **Effects of model-based correction on transcriptome-wide coraltaion between decidua to fetal expression ratio (D:F) and proportion of the maternally inherited allele in the fetal placenta**. **A:** Dotplot shows the averaged expression proportions of the PWK allele in the y axis and log transformed D:F in fetal placenta for all genes in samples from ♀mus$^{PWK}$ x ♂mus$^{WSB}$ crosses before correction. **B:** Same plot generated after correcting for maternal contamination.

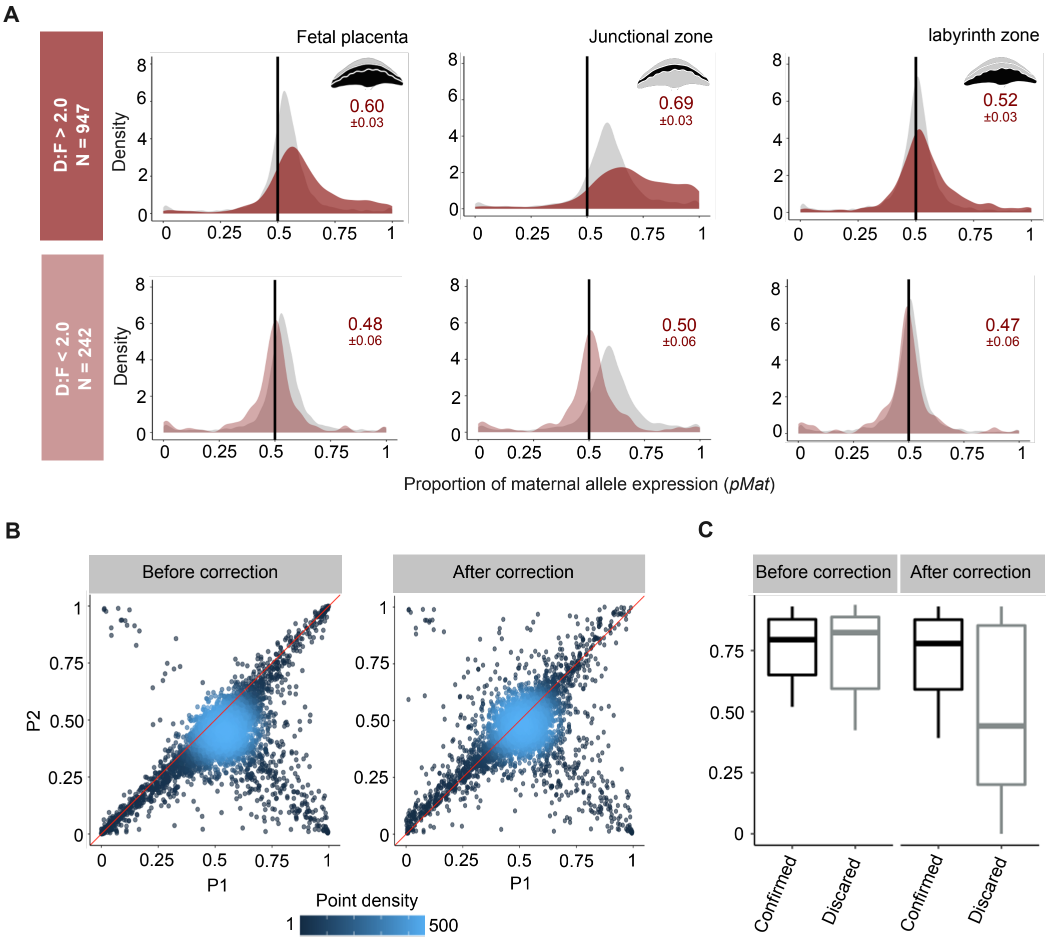


Figure 3: **Effects of maternal contamination across fetal placental layers and effects of model-based correction**. **A:** Density plots show in red the distribution of LEW alleles (maternal) expression ratio across two subsets of genes in fetal tissues sampled from ♀mus$^{LEW}$ ♂mus$^{WSB}$ crosses. Vertical black line in x axis is placed at 0.5 for reference. Gray histograms show transcriptome-wide distributions for each layer. Dark red histograms (top) show distribution across genes with a high maternal to fetal expression ratio (D:F ¿ 2, N = 947 genes); Light red histograms (bottom) show distributions across genes with a low maternal to fetal expression ratio (D:F ¡ 0.2, N = 242 genes). Numbers on each histogram show the center of the distribution of target gene sets followed by a 95% confidence interval estimated from from a binomial model. **B:** Genome wide distribution of P1 and P2 values estimated from fetal placenta samples of the same cross before and after correcting for maternal contamination. **C:** Boxplots show the distribution of maternal bias scores of 19 genes scrutinized in Okae et. al (2012) before and after correcting for maternal contamination using our model-based correction. Genes are grouped in two categories: black box corresponds to genes with confirmed imprinted expression and grey box contains genes discarded as false positives in Okae et al (2012).

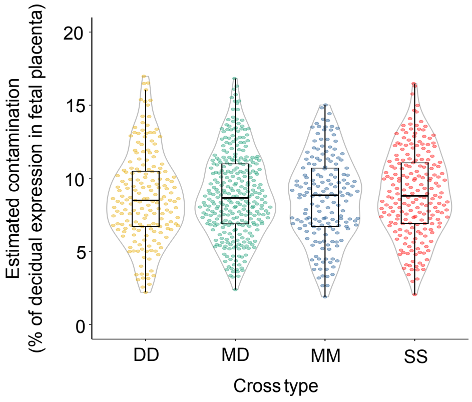


Figure 4: **Estimates of maternal contamination**. Box plots show the distribution of estimates of maternal contamination calculated as the percent of decidual expression required to explain contaminatino in the fetal placenta. Each boxplot displays estimates generated from diagnostic gene sets from the four cross types in this study. Cross types are coded as: **DD:** dom x dom, **MD:** mus x dom, **DD:** dom x dom, and **SS:** spret x spret.

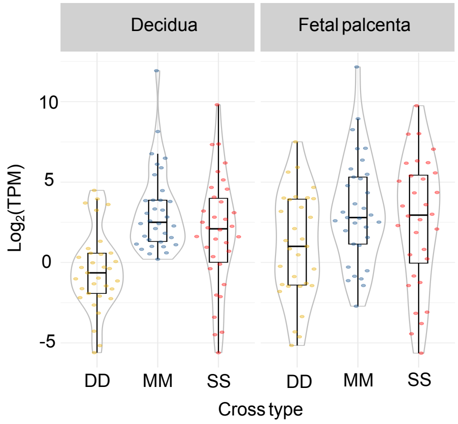


Figure 5: **Increased differential expression in the decidua in a subset of co-regulated genes**. Violin plots show standardized average expression levels in decidua and feal palcenta samples of 34 genes clustered in a WGCNA module enriched for maternally biased expression (module G) and showing divergent expression levels in *M. m. domesticus*. Cross types are coded as: **DD:** dom x dom, **MD:** mus x dom, **DD:** dom x dom, and **SS:** spret x spret.

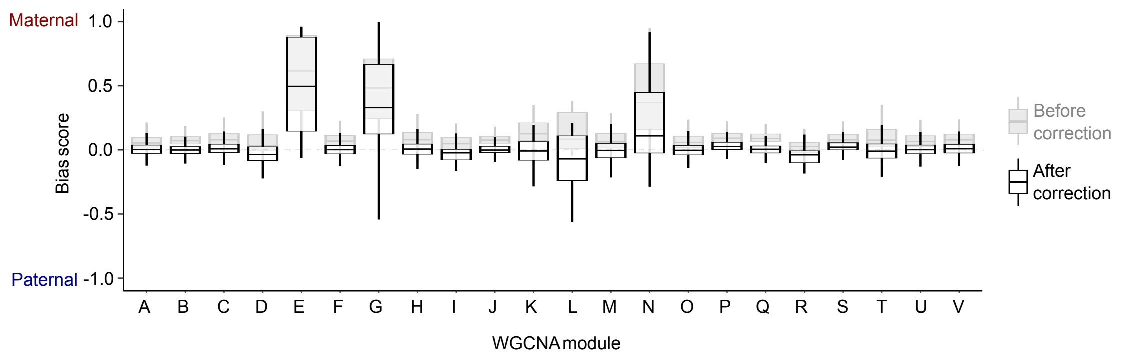


Figure 6: **Changes in the distribution of bias scores across WGCNA modules after correcting for contamination**. Background boxplots (gray) show the distribution of bias scores across WGCNA modules before correction. Boxplots with black lines show same distributions after correction. Note the largest shift towards biallelic expression in module "N" suggesting most bias at this module is laregely explained by contamination.

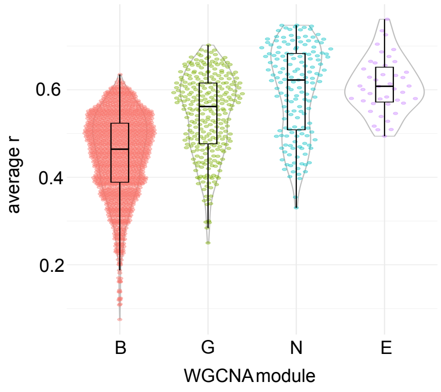


Figure 7: **Connectivity of maternally biased modules**. Violin plots show the average intramodular connectivity of four WGCNA modules showing enrichment for maternally biased expression. Intramodular connectivity is a metric of the strength of covariation in expression levels among genes in the module and higher numbers provide greater support of coregulation.

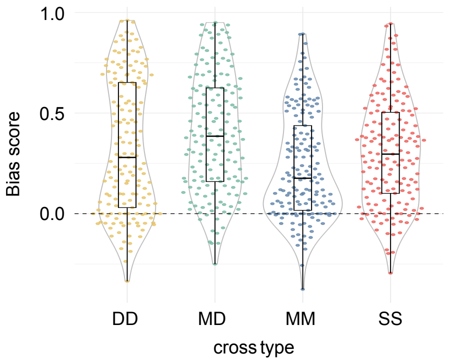


Figure 8: **Differences in the extent of maternal bias across cross types**. Violin plots show the distribution of bias scores among genes clustered in the "N" WGCNA module after correcting for maternal contamination. Cross types are coded as: **DD:** dom x dom, **MD:** mus x dom, **DD:** dom x dom, and **SS:** spret x spret.
