## Supplementary material for "Evolution of parent-of-origin effects on placental gene expression in house mice": Suplemental methods

### Model based correction of maternal contamination

Maternal and embryonic tissues are tightly integrated in the mouse placenta resulting in inevitable contributions of maternal cells to estimates of gene expression in dissected fetal placenta samples (Proudhon and Bourc'his 2010). Maternal contamination in the mouse placenta has been modeled in the past (Finn et al. 2014), however contamination estimates varied dramatically depending on the set of diagnostic genes used. Below, we describe the strategy we used to measure maternal contamination using a modification of that implemented by Finn et al. (2014) aimed to increase the accuracy of contamination estimates.

*Model of allelic expression in fetal placenta samples*

Our model assumes that maternal contamination originates from remnant decidual cells in dissected fetal placenta samples. Thus, we modeled gene expression in the fetal placenta ($exp_{Total}$) as the sum of transcripts originating from the maternally inherited allele ($exp_{Mat}$), the paternally inherited allele ($exp_{Pat}$) and maternal alleles in contaminating decidual cells ($exp_{Cont}$)(equation 1).

$$exp_{Total} = exp_{Mat} + exp_{Pat} + exp_{Cont} \qquad (1)$$

Because inbred genetic lines were used for experimental mice crosses, dams are expected to be homozygous for most loci in the genome, thus, maternal transcripts from decidual cells cannot be differentiated from transcripts originating from the maternally inherited allele in fetal placenta cells using sequencing data. Previous efforts to measure contamination used loci known to display complete silencing of the maternal allele in the fetal placenta (*i.e., expMat* = 0), where maternal expression can be assumed to originate exclusively from contaminating cells and $exp_{Cont}$ can be estimated as the difference between $exp_{Total}$ and $exp_{Pat}$ (equation 2). However, these genes represent a small sample of the transcriptome, (N=10) and did not yield consistent estimates of contamination, presumably due to variation in the extent of maternal silencing and inconsistent levels of expression across replicates. To overcome this limitation we used a distinct set of genes as a diagnostic sample for measuring contamination.

*If $exp_{Mat} = 0$, then:*

$$exp_{Cont} = exp_{Total} - exp_{Pat} \qquad (2)$$

*Identification of diagnostic loci to measure maternal contamination*

Instead of using maternally silenced genes, we targeted genes with no allelic imbalance in the fetal placenta, where $exp_{Mat}$ and $exp_{Pat}$ can be assumed to be approximately equal, and $exp_{Cont}$ can be estimated as any excess from a 1:1 expression ratio of maternal and paternal alleles (equation 3).

*If $exp_{Mat} = exp_{Pat}$, then:*

$$exp_{Cont} = exp_{Total} - 2exp_{Pat} \qquad (3)$$

To identify a set of diagnostic genes, we generated a series of predictions on gene expression patterns consistent with no allelic imbalance in the fetal placenta:

1. *No differences in expression levels in decidua tissue between genetic lines used for a cross:* differences in gene expression levels between genetic lines could reflect *cis* regulatory variation of parental alleles, which would cause allelic imbalance in embryonic tissues. Thus, genes

showing no differential expression in the maternal decidua of both genetic lines used for a cross, were considered less likely to exhibit allelic imbalance in the fetal placenta.

2. *Maternal bias in the junctional zone and no bias in the labyrinth zone:* Our analysis on the transcriptome-wide effects of maternal contamination showed large effects on junctional zone tissue (which is directly juxtaposed to decidual tissue) and no effect on labyrinth zone tissue, therefore, a gene showing maternal bias in the junctional, but not the labyrinth zone, is likely to have no allelic imbalance and show maternal bias solely as a result of contamination.
3. *Allelic expression shifts towards maternal expression in the fetal placenta should be the same in the two directions of a reciprocal cross as long as the decidua to fetal expression ratio conserved between cross directions:* The extent of maternal bias in a gene is proportional to its decidua to fetal expression ratio (D:F). Therefore, we reasoned that genes with no allelic imbalance and with conserved D:F in the fetal placenta in both directions of a reciprocal cross should show consistent levels of maternal bias in both cross directions. We note that this might also be true for imprinted genes showing maternal bias, so we excluded known imprinted genes in the mouse fetal placenta. We also note that genes with allelic imbalance should not show this pattern given the asymmetry of true allelic proportions between reciprocal crosses (figure 1).

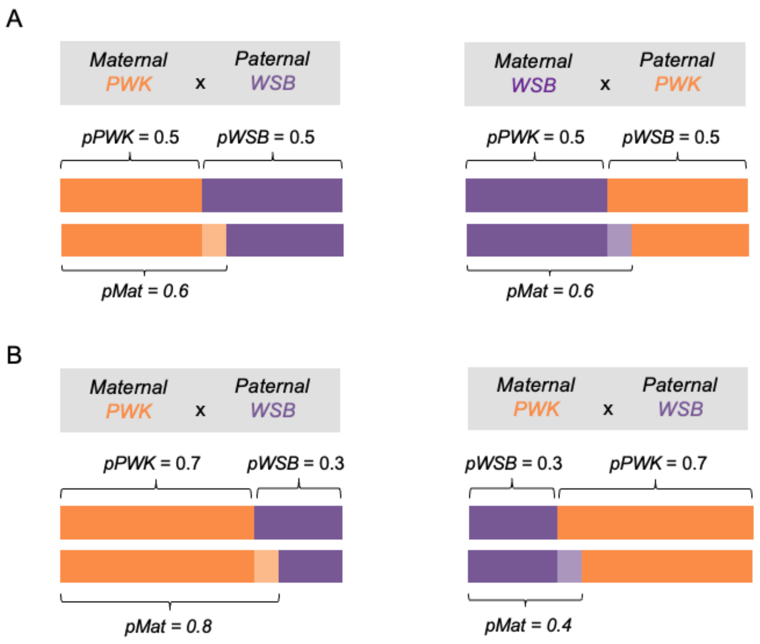


**Figure 1. Contamination on genes without allelic imbalance results in equal estimates of *pMat* in both directions of a reciprocal.** The schemes show theoretical allele frequencies with and without contamination in both directions of a reciprocal cross (♀PWK x ♂WSB and ♀WSB x ♂PWK) in a gene without allelic imbalance (A) and one with allelic imbalance (B). Solid bars represent the contribution of parentally inherited alleles in embryonic tissue and light color bars represent the effect of contamination on estimates of allelic expression ratios assuming a +0.1 shift of the maternal allele expression ratio induced by contamination. **A)** When a gene has no allelic imbalance (*pPWK*=0.5, *pWSB*=0.5), a shift of +0.1 in maternal allelic expression results in the same value of *pMat* (0.6) in both directions of the cross **B)** In contrast, when a gene shows allelic imbalance (*pPWK*=0.7, *pWSB*=0.3) estimates of *pMat* differ between the two directions of the cross due to the asymmetry in the true expression ratio of the maternal allele.

We identified genes fitting all three predictions in each cross type separately, and used them as diagnostic genes to measure maternal contamination in the fetal placenta. To avoid noise incorporated by sampling error on expression and allelic ratios, we filtered our diagnostic gene set excluding genes with low expression levels (TPM < 2) in either the decidua or the fetal placenta and genes with low power for assessing allele specific expression (*i.e* less than 2 diagnostic variants between genetic lines).

*Calculating average contamination*

The size of diagnostic gene sets, varied by cross type ranging from 169 to 308 with the largest number of diagnostic genes in the cross between the most divergent lines as expected from increased power for allele specific expression. We used these genes to estimate $exp_{Cont}$ using equation 3 and calculated contamination ($C$) as the average proportion of a gene's expression level in the decidua ($exp_{Dec}$) required to explain maternal contamination in the fetal placenta (equation 4). This value should be consistent across genes and determined by the amount of remnant decidual tissue in a fetal placenta sample.

$$C = \frac{1}{N} \sum_{i=1}^{N} \frac{exp_{Cont_i}}{exp_{Dec_i}} \quad (4)$$

*Accounting for contamination and re-calculating P1 and P2*

We calculated the theoretical value of $exp_{Cont}$ for each gene in the transcriptome as a function of its expression level in the decidua and the estimated average contamination (equation 5). Next, we accounted for contamination in the transcriptome by subtracting $exp_{Cont}$ from the observed levels of maternal ($O.exp_{Mat}$) and total expression ($O.exp_{Total}$) in the fetal placenta and re-calculated the P1 and P2 values used to estimate parent-of-origin bias score (equations 6 and 7).

$$exp_{Cont} = exp_{Dec} \times C \quad (5)$$

$$P1_{adj} = \frac{O.exp_{Mat} - (exp_{Dec} \times C)}{O.exp_{Total} - (exp_{Dec} \times C)} \quad (6)$$

$$P2_{adj} = \frac{exp_{Pat}}{O.exp_{Total} - (exp_{Dec} \times C)} \quad (7)$$

To account for variation in decidual expression levels between genetic lines, estimates of decidual expression were generated individually for each maternal line. All expression levels were estimated from RNA-seq data and standardized using transcripts per million reads (TPM).
